# Genome-wide Screens Identify Lineage- and Tumor Specific-Genes Modulating MHC-I and MHC-II Immunosurveillance in Human Lymphomas

**DOI:** 10.1101/2020.03.13.989558

**Authors:** Devin Dersh, James D. Phelan, Megan E. Gumina, Boya Wang, Jesse H. Arbuckle, Jaroslav Holly, Rigel J. Kishton, Tovah E. Markowitz, Mina O. Seedhom, Nathan Fridlyand, George W. Wright, Da Wei Huang, Michele Ceribelli, Craig J. Thomas, Justin B. Lack, Nicholas P. Restifo, Thomas M. Kristie, Louis M. Staudt, Jonathan W. Yewdell

## Abstract

Tumors frequently subvert MHC class I (MHC-I) peptide presentation to evade CD8+ T cell immunosurveillance. To better define the regulatory networks controlling antigen presentation, we employed genome-wide screening in human diffuse large B cell lymphomas (DLBCLs). This approach revealed dozens of novel genes that positively and negatively modulate MHC-I cell surface levels. Identified genes cluster in multiple pathways including cytokine signaling, mRNA processing, endosomal trafficking, and protein metabolism. Many genes exhibit lymphoma subtype- or tumor-specific MHC-I regulation, and a majority of primary DLBCL tumors display genetic alterations in multiple regulators. We establish that the HSP90 co-chaperone SUGT1 is a major positive regulator of both MHC-I and MHC-II cell surface expression. Further, pharmacological inhibition of two negative regulators of antigen presentation, EZH2 and thymidylate synthase, enhances DLBCL MHC-I presentation. These and other genes represent potential targets for manipulating MHC-I immunosurveillance in cancers, infectious diseases, and autoimmunity.

## Introduction

CD8+ T cells surveil for malignant or pathogen-infected cells by probing MHC class I (MHC-I), a cell surface heterotrimer constitutively expressed on virtually all nucleated cells in jawed vertebrates. MHC-I complexes are formed in the endoplasmic reticulum (ER) by association of polymorphic type I integral membrane proteins with β_2_m light chains, subsequently loaded with high affinity peptides transported from the cytoplasm by the peptide transporter TAP. Peptide-loaded MHC-I is trafficked through the Golgi complex to the plasma membrane where it can be scanned by T cells via their clonally restricted T cell receptors (TCRs), which are selected for their ability to recognize self MHC heavy chains bound to peptides perceived as non-self.

Debate raged for decades regarding the ability of T cells to recognize cancer cells as non-self. The remarkable success of T cell-based immunotherapy has ended the debate, clearly establishing the relevance of pioneering work defining tumor-specific peptides in mouse models (Van den Eynde et al., 1995; van der Bruggen et al., 1991). Immunogenic cancer-associated peptides can arise from mutations in tumor cell source genes (Yadav et al., 2014), cellular alterations that enhance transcription or translation of non-tolerized open reading frames (Laumont et al., 2018), and post-translational modifications of peptides (Liepe et al., 2019).

One of the most compelling findings supporting the clinical potential of immunotherapy was the relatively facile identification of cancer cells that downregulate or inactivate components of the antigen processing and presentation (APP) pathway (Esteban et al., 1990; Hellstrom, 1960; Tran et al., 2016). Since such immunoevasion is one of the principal reasons for immunotherapy failure (Sade-Feldman et al., 2017), it is critical to identify gene products that enable or counteract immunoevasion.

To study cancer cell antigen presentation from an unbiased perspective, we conducted forward genetic screens for regulators of MHC-I in diffuse large B cell lymphoma (DLBCL). DLBCL is the most common adult lymphoid malignancy, the most frequent form of non-Hodgkin lymphoma, and a significant public health burden. With frequent mutations, common localization in lymph nodes (a primary site of T cell activation), and constitutive expression of both MHC-I and MHC-II, DLBCLs are likely subjected to high levels of immunosurveillance. Indeed, immune evasion is common in DLBCL, with an estimated ∼40-75% of biopsies showing aberrant MHC-I expression or localization (Challa-Malladi et al., 2011; Ennishi et al., 2019; Nijland et al., 2017), though genetic analyses have thus far been unable to explain all MHC-I^low^ patient samples.

Rational deployment of immunotherapy will ultimately require deeper understanding of tumor-specific genetic variation, both in antigen presentation and in oncogenic signaling in general. In DLBCL, for example, genetic analyses of different tumors combined with survival trends from standard immunochemotherapies identified clear genetic subtypes (Alizadeh et al., 2000; Schmitz et al., 2018). The “cell of origin” model categorizes tumors based on the differentiation state of their origin B cell. Activated B cell-like (ABC) DLBCL shows gene expression patterns similar to that of activated peripheral B cells and is clinically aggressive. Germinal center B cell-like (GCB) DLBCL likely derives from B cell centrocytes and is more responsive to standard immunochemotherapy (Read et al., 2014; Rosenwald et al., 2002). Even cells of the same classification can be widely heterogenous, highlighting a need for deeper personalization in treatment plans, including immunotherapies.

In the present study, we used genome-wide CRISPR screening to investigate the regulation of cell surface MHC-I expression in both ABC and GCB DLBCL patient-derived tumor cells. We identify and validate dozens of novel regulators of antigen presentation, demonstrating remarkable coordination of multiple cellular pathways. Our findings show that control of antigen presentation and immunoevasion varies between tumor lineages and even tumors of the same lineage. We describe examples of cellular complexes that can be targeted to enhance tumor cell immunogenicity and improve immunotherapies. The work represents the most comprehensive global analysis of antigen presentation regulation to date and points to unique modes of regulation that may be utilized in diverse cancers.

## Results

### Genome-wide CRISPR/Cas9 screens identify regulators of MHC-I in DLBCL

We identified genes that regulate surface MHC-I in DLBCL using four patient-derived lymphoma cell lines representing both GCB (SUDHL5, BJAB) and ABC (HBL1, TMD8) classified tumors. Cells were stably transduced with a doxycycline-inducible Cas9 cassette (Phelan et al., 2018) followed by transduction with the Brunello genome-wide CRISPR knockout (KO) library (Doench et al., 2016) (screen outline depicted in Figure 1A). Through two rounds of flow cytometry-based sorting, we separated populations with low and high MHC-I surface expression using the pan human MHC-I specific W6/32 monoclonal antibody and sequenced resulting genomic DNA to determine sgRNA copy numbers (Figure 1B). This multi-sort strategy was critical in filtering variation inherent in MHC-I expression in normal distributions of cell populations. For example, MHC-I levels of sorted cells changed considerably after just a week of growth post-sorting (Figure 1B).

**Figure 1.**
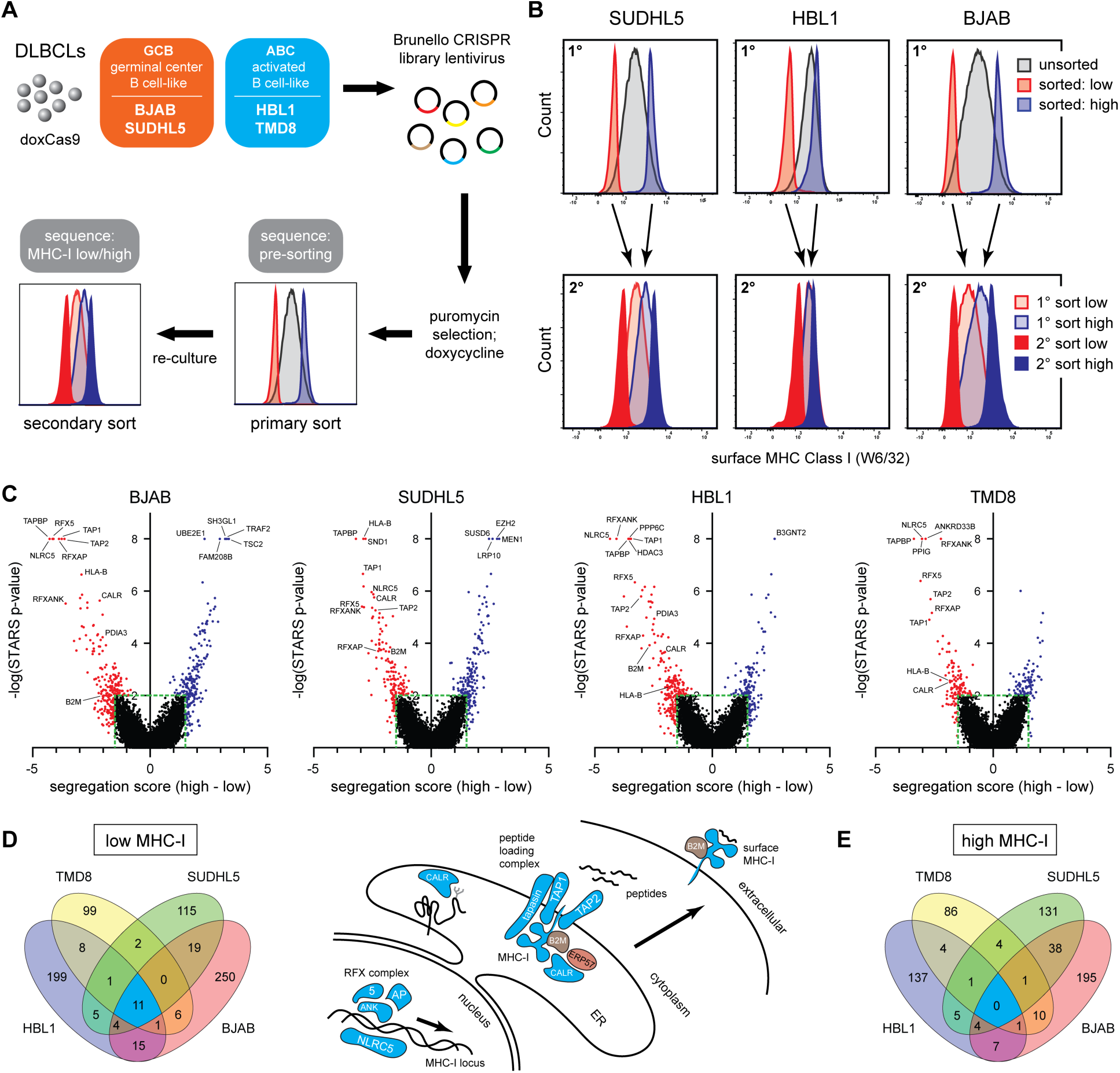
MHC-I genome-wide screen setup, sorting, and analysis. **(A)** Overview of the CRISPR/Cas9 screens used to identify regulators of MHC-I in diffuse large B-cell lymphoma tumor lines. **(B)** Example flow cytometry histograms of the sorting for MHC-I low/high cells using the pan-MHC-I antibody W6/32 (1°, primary sort; 2°, secondary sort). **(C)** STARS statistical analysis of each gene plotted against segregation scores in the sorted populations relative to input controls. Negative segregation score: sgRNAs enriched in MHC-I^low^ population; positive segregation score: sgRNAs enriched in MHC-I^high^ population. Red/blue indicate genes with p-value of < 0.01 or a segregation score outside the range of −1.5 – 1.5. Note that some genes were set at -log(p-value) of 8 due to p-value rounding in STARS. **(D)** (left) Venn diagram of top gene KOs from the MHC-I^low^ analyses in four tumor lines. (right) Schematic depicting some genes known in antigen processing and presentation and their identifications in the CRISPR screens – coloring refers to Venn diagram overlap colors. **(E)** Venn diagram of top gene KOs from the MHC-I^high^ analyses.

Counts of individual Brunello sgRNA sequences were used to calculate gene-specific segregation scores – Z-score-based measurements of the ratio of sgRNA sequences in MHC-I high vs. low populations. STARS software was also used to score genes through ranks and numbers of enriched sgRNAs in sorted populations relative to input controls (Doench et al., 2016) (Figure 1C; Supplemental Table 1).

Importantly, nearly all genes previously described as APP regulators were highly ranked in our analysis, including all components of the peptide loading complex and known MHC-I transcription factors (Figures 1C and 1D). We identified dozens of putative novel positive and negative regulators – some specific for a particular tumor and others shared between two or more tumors (Figures 1D and 1E; Supplemental Table 2).

### Validation and pathway analysis of screen hits

Although powerful, phenotypic CRISPR screens typically generate significant numbers of false positives and negatives. To validate positive MHC-I regulators, we cloned sgRNAs for each of 130 potential hits revealed by the genetic screens. These were generally selected based on high STARS ranking but also included lower scoring hits that clustered into clear functional complexes via STRING analysis (Szklarczyk et al., 2019). Cells were infected with lentiviruses containing targeting sgRNAs and an EGFP expression cassette, or non-targeting sgRNAs without EGFP expression, and cultured identically. Surface MHC-I levels were measured 9-11 days post-infection. For each KO line, edited and control-infected cells were mixed and analyzed simultaneously, using EGFP expression to distinguish edited cells (Figure 2A). At the same time, we measured expression of surface MHC class II (HLA-DR) as well as CD147, which is not known to participate in antigen presentation and therefore controls for non-APP-specific effects on cell surface protein biogenesis or turnover.

**Figure 2.**
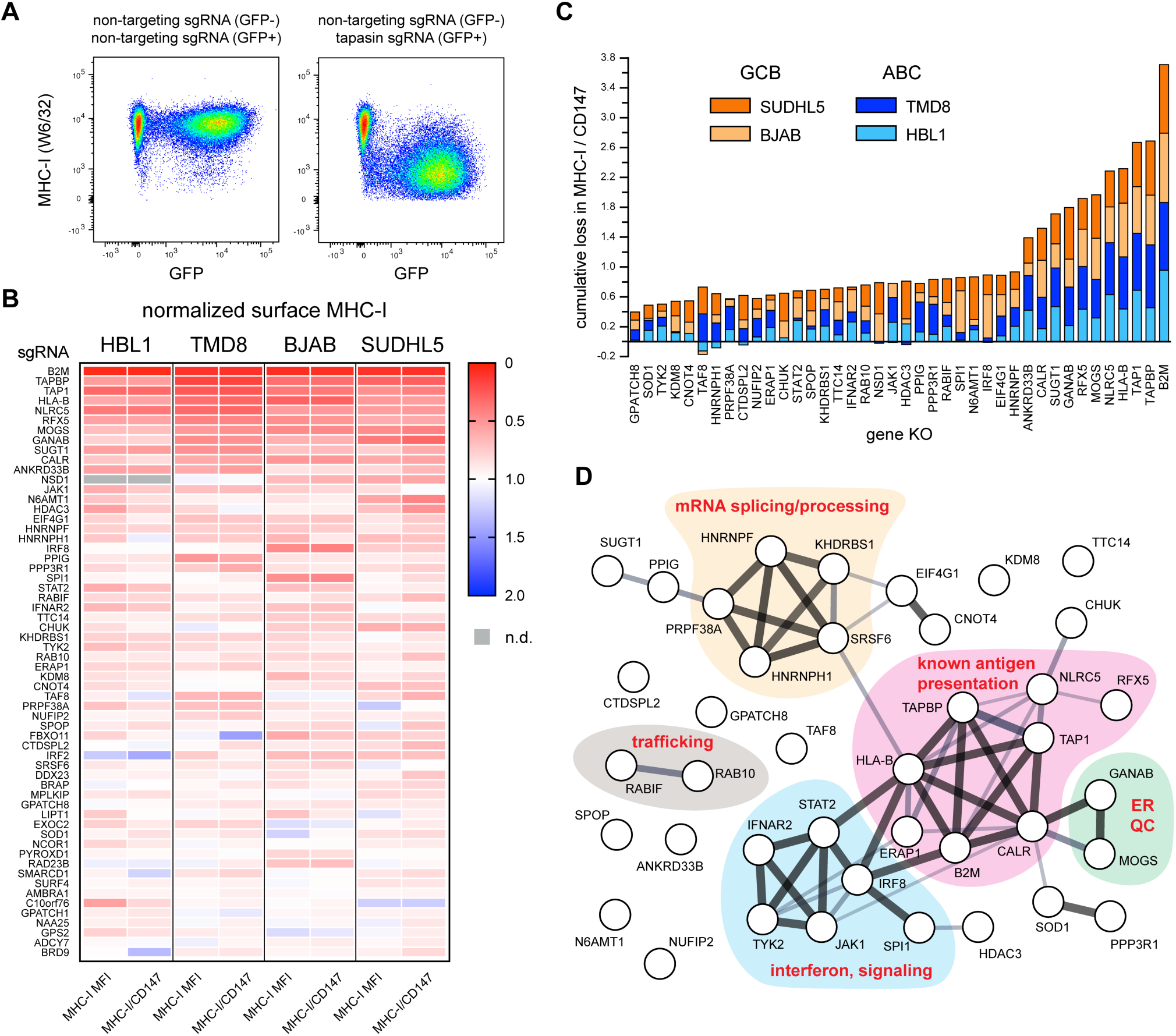
Validation of positive regulators of MHC-I. **(A)** Example of how individual gene KOs were tested for surface MHC-I effects. Cells were independently infected with lentiviruses encoding targeting or non-targeting sgRNAs in GFP+ or GFP- backbones. After 8-11 days of selection and Cas9 induction, cells were mixed, stained, and analyzed together by flow cytometry. GFP- and GFP+ populations were used to quantify remaining surface MHC-I via the pan-MHC-I antibody W6/32. **(B)** Validations of positive regulators of MHC-I. 60 of the top performing gene KOs across four parental tumor lines are displayed as a heatmap of surface MHC-I levels relative to non-targeting sgRNA. Also shown are quantifications of MHC-I per CD147, an unrelated surface protein, to highlight global protein trafficking alterations. Note that not every previously known regulator was reanalyzed (e.g., TAP2, RFXAP). **(C)** Cumulative loss in MHC-I/CD147 upon indicated gene KO across the four tumor lines. GCBs orange; ABCs blue. **(D)** STRING analysis of the validated positive regulators of MHC-I in DLBCL; thickness of line indicates relative confidence in genetic interaction.

We used the positive regulator validation panel to test each of the four DLBCL cell lines from the CRISPR screens (top 60 validated genes shown in Figure 2B; see also Supplemental Table 3). Dozens of gene KOs showed consistent loss of surface MHC-I across the different lymphoma models (Figure 2C). These genes play roles in MHC-I antigen presentation, ER quality control, interferon signaling, trafficking, and RNA processing and splicing, among other functions (Figure 2D).

We similarly validated predicted APP negative regulators using a panel of 70 individual sgRNAs based on top screen hits, the majority of which were confirmed (Figure 3A; Supplemental Table 3). Repressive genes were particularly employed by GCB tumors relative to ABC cells (Figure 3B, orange vs blue). The negative regulators cluster into several functional pathways that include mTOR regulation, mRNA capping and translation, the PRC2 repressor complex, components of the ubiquitin/proteasome system, as well as numerous endolysosomal trafficking factors likely important for the internalization of MHC-I and its endosomal recycling and/or lysosomal degradation (Figure 3C).

**Figure 3.**
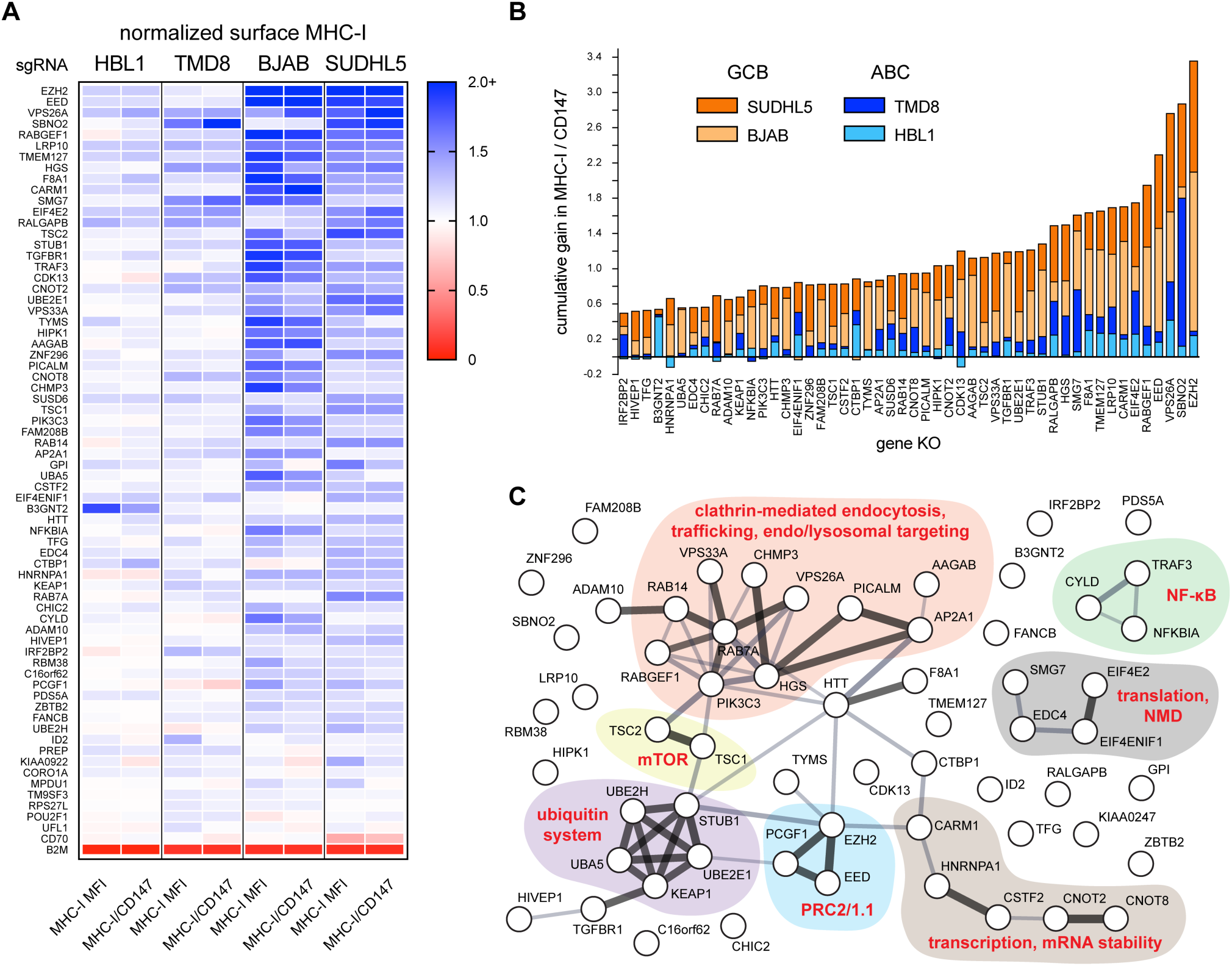
Validation of negative regulators of MHC-I. **(A)** Validations of negative regulators of MHC-I. 69 gene KOs across four parental tumor lines are displayed as a heatmap of surface MHC-I levels relative to non-targeting sgRNA. B2M KO was also included as a control for Cas9 activity and antibody staining. Also shown are quantifications of MHC-I per CD147, an unrelated surface protein, to highlight global protein trafficking alterations. **(B)** Cumulative gain in MHC-I/CD147 upon indicated gene KO across the four tumor lines. GCBs orange; ABCs blue. **(C)** STRING analysis of the validated negative regulators of MHC-I in DLBCL; thickness of line indicates relative confidence in genetic interaction.

MHC-I polymorphism potentially contributes to variability in the effects of a gene KO for a given tumor (Supplemental Figure 1). We did not, however, observe major differences in the behavior of HLA-A2 in HBL1 cells compared to total MHC-I levels across a variety of genetic manipulations (Supplemental Figure 2A). Separately, we observed that both segregation score (Supplemental Figure 2B) and STARS significance values (Supplemental Figure 2C) from the CRISPR screens correlated with the magnitude of MHC-I alteration in subsequent validation experiments, confirming the power of the genetic screens.

While most regulators of APP affected MHC-I surface levels similarly across multiple tumors, examples of lineage or tumor specificity were also clear (Supplemental Figure 3 summarizes all KOs in all lines). For instance, ABC tumors were much more resistant to enhancement of surface MHC-I (see Figure 5). We even identified genes whose deletion exerted *opposite* effects on MHC-I expression in different tumor lines (Figure 4A), highlighting the genetic heterogeneity so common within DLBCL (Schmitz et al., 2018). The opposing effects of BCOR and BAP1 regulation of MHC-I in BJAB and SUDHL5 cells are particularly surprising, given that both lines are classified as GCB tumors and share relatively similar gene expression profiles.

**Figure 4.**
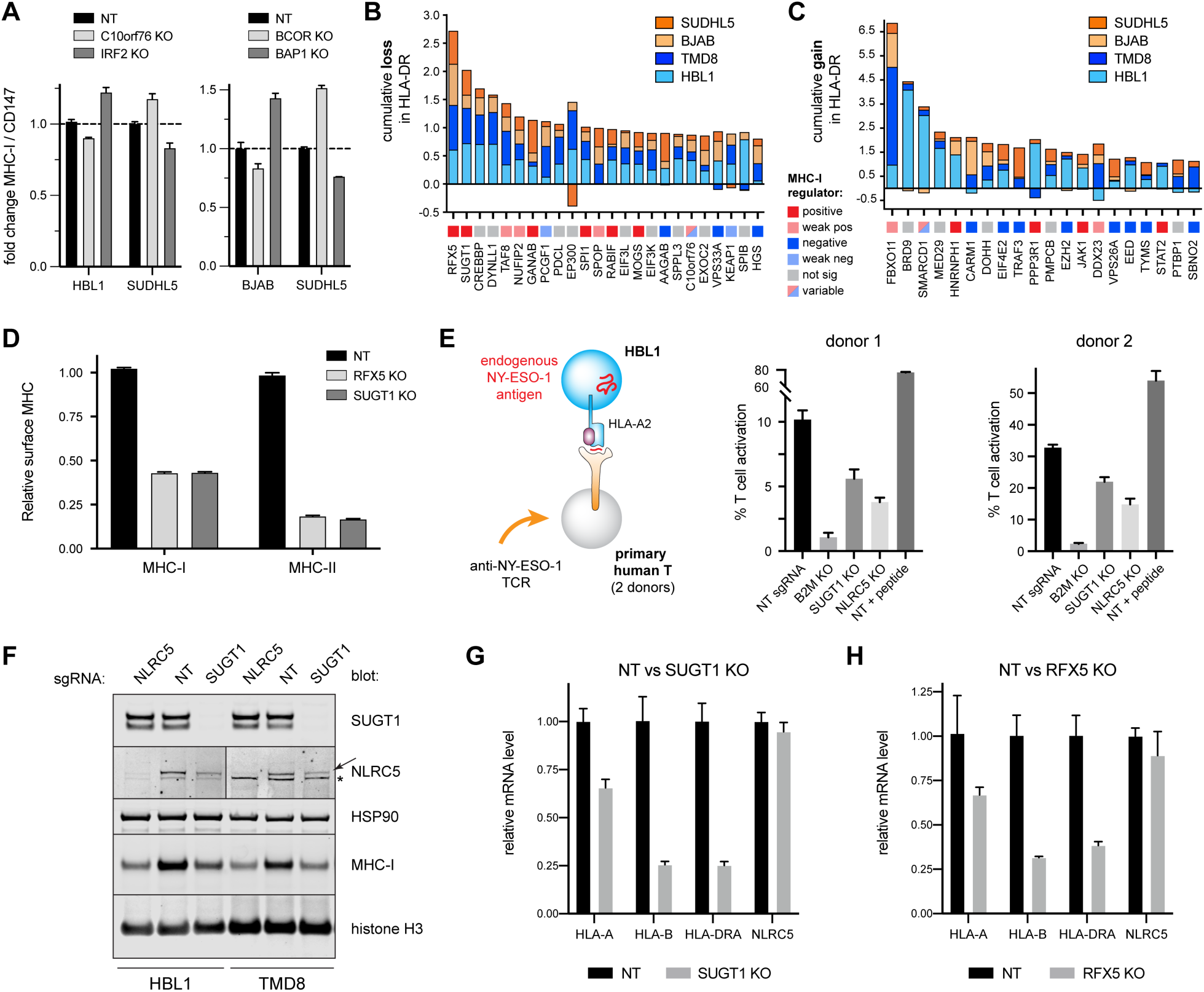
Specificity of MHC regulation and the role of SUGT1 in antigen presentation. **(A)** Opposing effects of gene KOs on surface MHC-I levels of different tumor lines as measured by flow cytometry. **(B)** Top performing gene KOs for the loss of surface HLA-DR, quantified across different tumor cells. GCBs orange; ABCs blue. Each gene is also classified with its MHC-I regulator status (bottom boxes). **(C)** Same as **B**, but for cumulative gains in HLA-DR upon gene KO. **(D)** Effects of RFX5 or SUGT1 KO on the surface levels of MHC-I and MHC-II in TMD8. **(E)** (left) Schematic of T cell co-culture assay. HBL1 cells, which are HLA-A2^+^ and natively express the cancer testis antigen NY-ESO-1, were co-cultured with primary human T-cells transduced with a TCR recognizing the NY-ESO-1 peptide 157-165 restricted by HLA-A2. (right) T-cells were monitored for activation by 4-1BB upregulation after 12-14 hours with the indicated HBL1 cells. NT, non-targeting sgRNA. NT + peptide, non-targeting sgRNA with exogenously added peptide, SLLMWITQV. T cells grown without target cells were manually set to 0% (1.14% average donor 1, 5.03% average donor 2). **(F)** NLRC5 or SUGT1 were deleted in HBL1 or TMD8 cells, and whole cell lysates were subjected to Western blot analysis for the indicated proteins. Arrow, NLRC5. *, undetermined band from anti-NLRC5 antibody. **(G)** qPCR analysis of the indicated transcript levels in TMD8 cells modified with NT sgRNA or SUGT1 KO. **(H)** Same as **G**, but with RFX5 KO. For entire figure, bar graphs represent mean with standard deviations, minimum n = 3.

**Figure 5.**
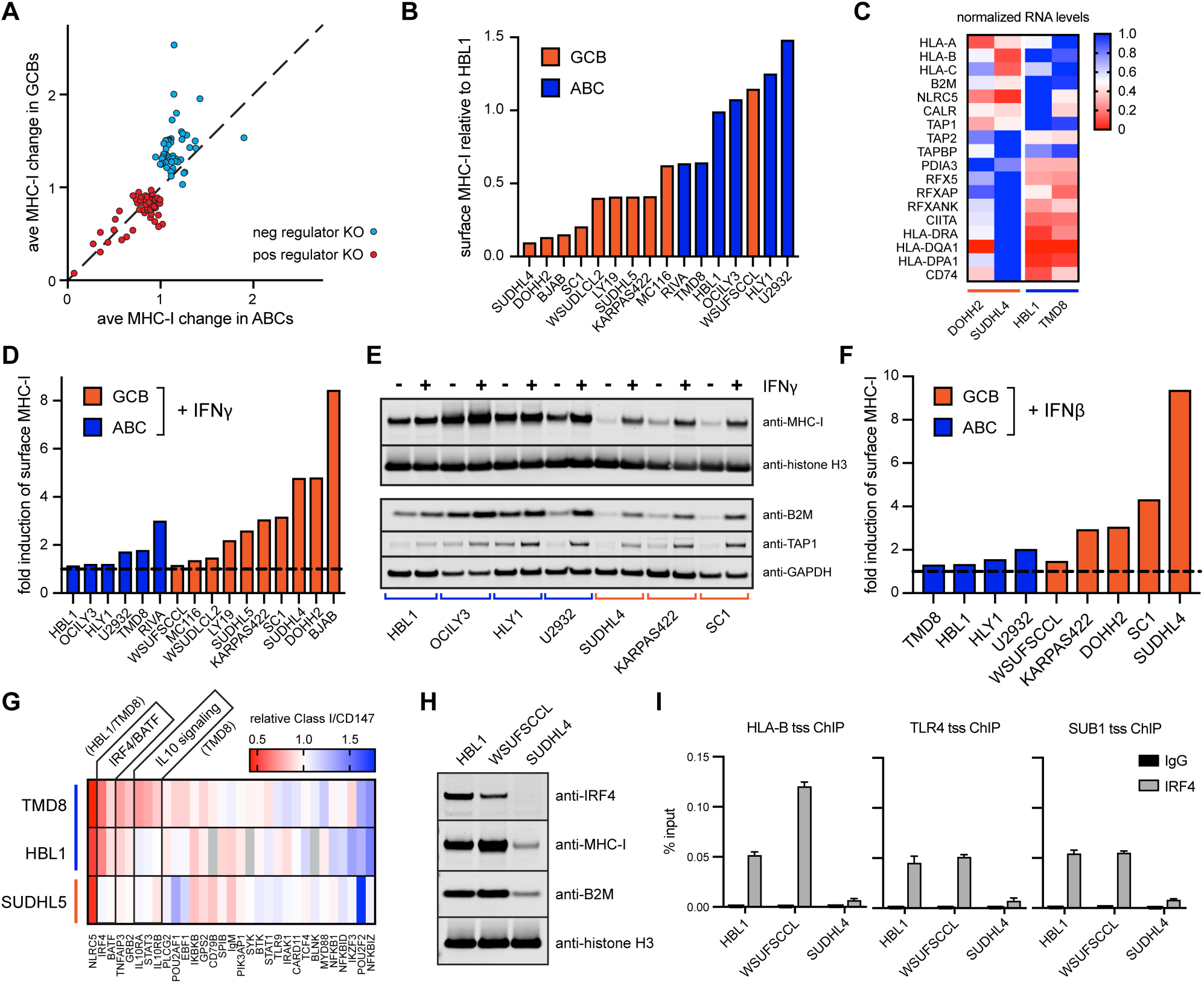
ABC DLBCLs drive antigen presentation. **(A)** The fold change in surface MHC-I of a given gene KO is plotted as an average between GCBs (SUDHL5, BJAB) and between ABCs (HBL1, TMD8). The top 50 positive regulators and top 50 negative regulators are displayed. Diagonal line indicates an equivalent response between GCBs and ABCs. **(B)** A panel of 16 DLBCL lines were measured for surface MHC-I by flow cytometry, and MFI were normalized to WT HBL1 cells. GCBs orange; ABCs blue. **(C)** Total RNA was isolated from the indicated cells and subjected to RNAseq analysis. Each transcript is normalized to the highest FPKM value of the four lines. **(D)** The indicated cells were treated for 2 days with 0 or 500U/mL IFNγ; fold induction of surface MHC-I with treatment is plotted. **(E)** Cells were left untreated or treated with 500U/mL IFNγ followed by Western blot analysis of whole cell lysates. **(F)** The indicated cells were treated for 2 days with 0 or 500U/mL IFNβ; fold induction of surface MHC-I with treatment is plotted. **(G)** TMD8 (ABC), HBL1 (ABC), and SUDHL5 (GCB) cells were transduced with sgRNA for the indicated genes, and surface MHC-I complexes were measured compared to non-targeting controls after 7-8 days while cells retained viability. **(H)** HBL1, WSU-FSCCL, and SUDHL4 whole cell lysates were blotted with antibodies to the indicated proteins. **(I)** ChIP assays illustrate the occupancy of IRF4 at the transcriptional start sites (tss) of HLA-B, TLR4, and SUB1. qPCR was performed in triplicates, and data are displayed as means with standard deviations.

In conclusion, forward genetic CRISPR screens identified many known and novel regulators of MHC-I surface expression across multiple DLBCLs. Informed by these screens, we individually knocked out ∼1% of known human genes to measure the effects on antigen presentation and to validate *bona fide* regulators. This comprehensive genetic analysis of the MHC-I pathway highlights first, how multiple cellular processes coordinate in immunosurveillance, and second, that individual genes can participate in MHC-I function either globally or at a lineage- or tumor-specific level.

### Multiple genes co-regulate MHC-I and MHC-II

In individual validation experiments (Figures 2 and 3), we also examined the effect of gene KO on MHC-II cell surface levels, using a pan-HLA-DR specific mAb. B cells express high levels of cell surface MHC-II, the target of anti-tumor immunosurveillance mediated by CD4+ T cells, which can exhibit potent anti-tumor activity (Alspach et al., 2019).

Although the validation panel was selected for predicted effects on MHC-I, we identified a number of genes that regulated surface levels of HLA-DR, including RFX5, which is known to broadly participate in the transcription of both MHC-I and MHC-II loci (Meissner et al., 2012; Steimle et al., 1995), and CREBBP, which has been described to play a role in maintaining MHC-II expression in lymphomas (Figure 4B) (Green et al., 2015; Jiang et al., 2017). Interestingly, the identified regulators of MHC-II included a mix of genes which we identified as MHC-I positive regulators, MHC-I negative regulators, and some that did not show significant control over MHC-I (colored squares in Figures 4B and 4C).

Of these genes, we identified SUGT1 as a novel MHC-I and MHC-II co-regulator that mimics RFX5 function. In TMD8 cells, for example, SUGT1 and RFX KO had essentially identical effects, reducing surface expression of MHC-I by 50% and MHC-II by 80% (Figure 4D, p < 0.0001). To confirm the importance of SUGT1 knockdown on immunosurveillance, we utilized HBL1 cells, which endogenously express HLA-A2 and NY-ESO-1, a defined human cancer testis antigen (Robbins et al., 2015). SUGT1 KO in the tumor cells impaired activation of primary human T-cells expressing an HLA-A2-restricted TCR specific for the NY-ESO-1^157-165^ peptide (Figure 4E, p < 0.0001), confirming its requirement for T cell-mediated recognition of tumor cells.

The SUGT1 homolog in plants is an HSP90 co-chaperone required for the biogenesis of NLR gene family members (Mayor et al., 2007; Zhang et al., 2010). Could SUGT1 in DLBCLs assist in the folding of NLRC5 and CIITA, both members of the NLR family and master transcription factors, respectively, for MHC-I and MHC-II? Indeed, immunoblots show that SUGT1 KO decreases steady state levels of NLRC5 and total HLA-A/B/C by ∼50% (Figure 4F). Control over NLRC5 is post-transcriptional, as SUGT1 KO does not affect NLRC5 mRNA levels (Figure 4G, p = 0.2403 for NLRC5 mRNA, NT vs. SUGT1 KO). Loss of SUGT1 does, however, significantly reduce mRNA levels of HLA-A (p = 0.0017) and HLA-B (p = 0.0005), similar to the effect of losing the RFX5 transcription factor (Figures 4G and 4H). SUGT1 KO also significantly lowers HLA-DRA1 mRNA levels (Figure 4G, p = 0.0002), with a coordinate loss in MHC-II protein levels (Supplemental Figure 4). We did not observe a change in the steady state levels of CIITA in SUGT1-deleted cells by IP/Western, though it is possible that CIITA may be misfolded or hypofunctional.

Together, these data demonstrate that previously unidentified gene products such as SUGT1 make major contributions to MHC/peptide presentation, that genes can simultaneously co-regulate MHC-I and MHC-II (even in opposing directions), and that genetic alterations of these regulators can have major implications for T cell immunosurveillance of tumors.

### Oncogenic signaling enforces robust antigen presentation in activated B cell-like lymphomas

Our work validating regulators of MHC-I unveiled a clear cell-of-origin correlation: KO of MHC-I positive regulators had similar effects in GCB vs. ABC lines, whereas GCBs were much more responsive than ABCs to ablation of negative regulators (Figure 5A; see also cell-of-origin coloring in Figures 2C and 3B).

Importantly, the GCB lines tested express less overall surface MHC-I than the ABCs. Indeed, this pattern repeated in an expanded panel of 16 tumor lines (Figure 5B) and could not be attributed simply to increased cell surface area (Supplemental Figure 5A). In these analyses, we only included cells with measurable surface MHC-I and excluded tumor lines with mutations that completely prevented expression (e.g., B2M null).

Using RNAseq, we found that GCB tumors had lower levels of transcripts encoding class I heavy chains, β_2_m, TAP1, calreticulin, and NLRC5 than representative ABC tumors, and conversely had higher relative levels of MHC-II-associated mRNAs (Figure 5C). Thus, lineage-dependent MHC-I surface levels are likely controlled in part by transcription of APP genes. The high basal levels of antigen presentation in HBL1 and TMD8 also explain the relative difficulty in sorting ABC cells with significantly increased MHC-I during our genetic screens (see HBL1 in Figure 1B).

In contrast to the IFNγ- or IFNβ-mediated induction of MHC-I observed in most GCB cells, ABC lines were either totally unresponsive or only mildly responsive to type I and II interferons (Figures 5D-5F). This was not attributable to a lack of IFNγ receptors or the inability to activate the JAK/STAT signaling pathway (Supplemental Figures 5B-5D) and is consistent with the idea that ABC-specific signaling constitutively drives APP gene transcription to near saturation.

To identify factors in ABC tumors that contribute to high MHC-I gene expression, we knocked out a panel of 30 genes known to be critical for ABC DLBCL oncogenesis and signaling and measured MHC-I surface levels while cells retained viability (Figure 5G). KO of the IRF4/BATF complex substantially reduced surface MHC-I complexes in both HBL1 and TMD8 cells but not in a control GCB tumor line. IL10 signaling also clearly contributed to high antigen presentation in TMD8 but not HBL1 cells, highlighting heterogeneity between even similarly classified tumors.

IRF4/BATF are highly expressed in ABCs and required for cell survival (Yang et al., 2012); interestingly, they are also expressed in a subset of GCB tumors. For example, the GCB line WSU-FSCCL expresses IRF4, and this correlates with its high levels of MHC-I (Figures 5B and 5H). Via ChIP analysis, we observed that IRF4 associated with the HLA-B transcriptional start site in both an ABC line (HBL1) and the IRF4^+^ GCB WSU-FSCCL, but not in an IRF4^-^ GCB (SUDHL4) (Figure 5I). DNA enrichment was similar to that of genes known to be regulated by IRF4 (Yang et al., 2012). IRF4 peaks at HLA-B were also observed in published IRF4 ChIP-seq datasets of lymphoblastoid GM12878 (Gene Expression Omnibus, GSM803390; Supplemental Figure 5E).

Together, these findings indicate that oncogenic signaling inherent to ABC tumors can drive high expression of MHC-I pathway genes relative to GCB tumors. Such signaling is at least in part due to the activity of IRF4/BATF, which can associate directly with the HLA-B locus, the major source of classical MHC-I molecules in B cells. IRF4 may therefore be a pan-DLBCL predictor of immunogenicity, when considered in combination with other genetic alterations identified by our screens. High levels of APP genes and surface MHC-I molecules in ABC tumors have important genetic and clinical implications discussed below.

### Genetics of antigen presentation in patient tumors

APP genes are commonly altered in cancers, including DLBCL (Chapuy et al., 2018; Ennishi et al., 2019; Schmitz et al., 2018). We therefore examined the mutation status of the validated MHC-I regulators from our forward genetic screens in a cohort of 574 DLBCL patient biopsy samples.

Exome and RNA sequencing of DLBCL patient tumors confirmed the presence of mutations and genetic alterations in our list of validated positive MHC-I regulators (Figure 6A). While mutations occurred most frequently in MHC-I genes themselves (i.e. HLA-A/B/C and B2M), other now-validated MHC-I regulators showed significant genetic alteration frequency, implying clinical relevancy. Compared to GCB-type tumors, ABCs were clearly enriched for mutations in the canonical APP pathway.

**Figure 6.**
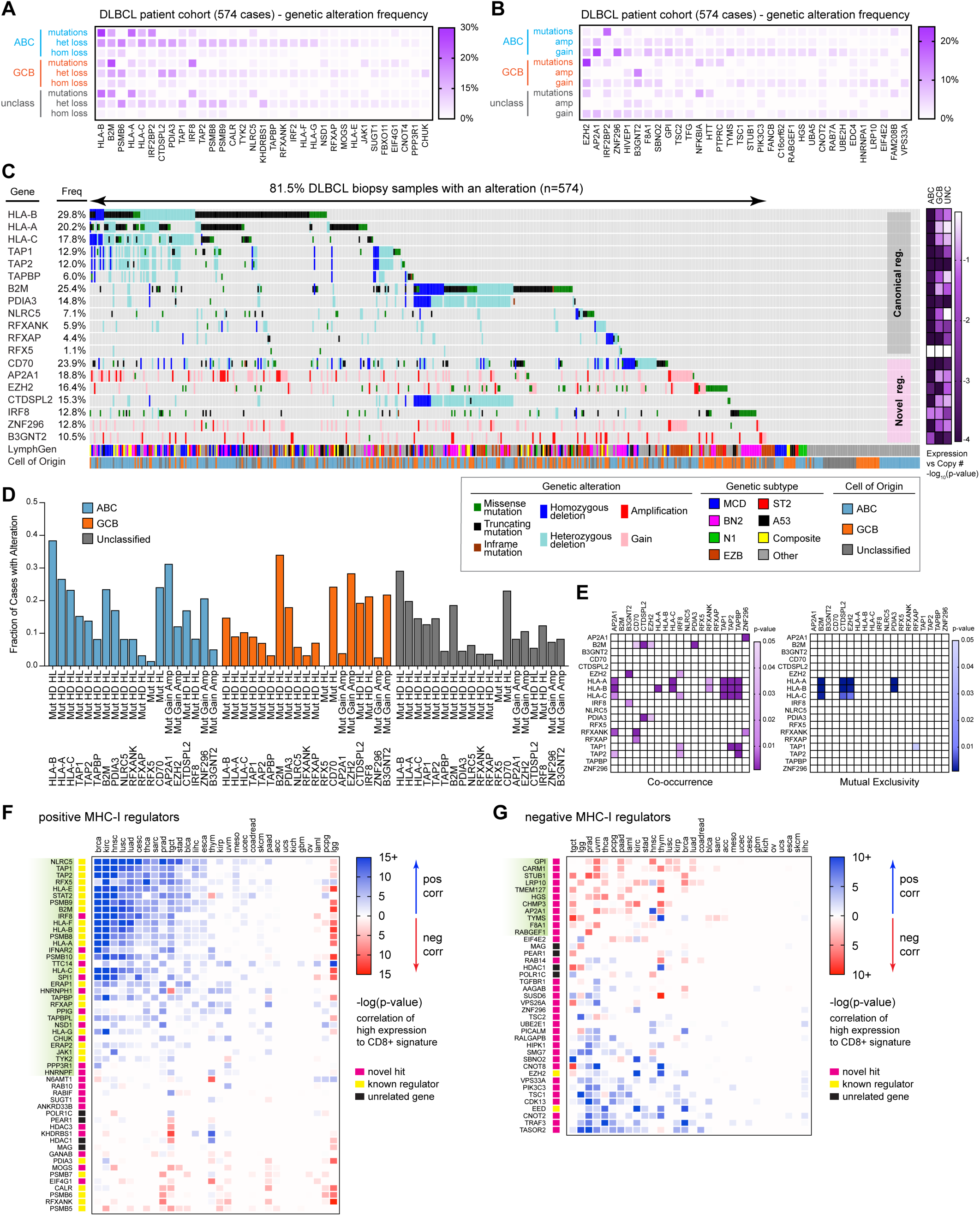
Genetic analysis of DLBCL cohort and pan-cancer correlations. **(A)** DLBCL biopsies (n=574) were analyzed by whole exome-seq, RNA-seq, and targeted amplicon deep sequencing (Schmitz et al., 2018). Validated MHC-I positive regulators are indicated with their frequency of genetic alterations. **(B)** Same as **A**, but with validated negative regulators of MHC-I. **(C)** Oncoprinter diagram of 574 DLBCL patient biopsy samples, indicating genetic alterations in each patient, their cell of origin (COO), and newest subtype classification (LymphGen, Wright et al., 2020). Right, -log(p-values) indicate likelihood of a non-zero slope from a linear regression of DNA copy number to gene expression for the indicated gene. **(D)** Frequency of genetic alterations in the indicated genes, separated by tumor COO categorization. Mut, mutation; HD, homozygous deletion; HL, heterozygous loss; gain, single copy gain; amp, multiple copy gain. **(E)** Co-occurrence and mutual exclusivity of genetic alterations in the indicated genes. **(F)** Using TCGA data, the indicated genes (positive regulators of MHC-I in DLBCL) were divided into the highest and lowest 33% of gene expression and correlated to CD8+ T cell signatures. Blue, positive T cell signature correlation with gene expression. Red, negative correlation. **(G)** Same as **F**, but for DLBCL negative regulator genes.

As with the positive MHC-I regulators, we also identified genetic mutations and chromosomal alterations in the validated negative regulators in DLBCL patient tumors (Figure 6B). In particular, GCB patients have a high likelihood of EZH2 gain-of-function mutations (Yap et al., 2011). Genetic gains in copy number were also commonly identified.

Limiting our analysis to those genes that showed a genetic alteration consistent with the screen phenotype (i.e. loss of a positive regulator, or gain of a negative regulator) and with allele frequencies >10%, we identified seven novel MHC-I regulators from our screen that were recurrently altered in DLBCL (Figure 6C). In total, ∼81% of DLBCL samples showed at least one form of genetic alteration among the list of class I regulators, though most tumors displayed hits in multiple genes. The identification of these newly validated genes may help explain previous findings of MHC-I negative DLBCL tumors that did not display characteristic mutations in classical MHC-I pathway genes (Challa-Malladi et al., 2011; Ennishi et al., 2019). The large fraction of tumors with mutations in MHC-I regulators strongly suggests that most DLBCL tumors are immunoedited by CD8+ T cell pressure. Importantly, copy number variations correlated with gene expression, implying the genetic losses and gains are functionally relevant (Supplemental Figure 6). Loss of heterozygosity in HLA genes is also a well-characterized immune evasion strategy separate from gene expression alterations (McGranahan et al., 2017), and individuals with HLA-B/C homozygosity are at enhanced risk for DLBCL (Wang et al., 2018). Additionally, genetic alterations were clearly focused around genes relevant to antigen presentation and were not only the byproduct of large deletion events (Supplemental Figures 7A and 7B).

Mutations in canonical class I pathway genes were recently characterized to be particularly common in some DLBCL subtypes (Schmitz et al., 2018). Our lineage analysis reveals that ABC tumors are most likely to have MHC-I pathway-related genetic alterations (Figure 6D). Of ABC tumors, the MCD subtype (MYD88^L265P^/CD79B mutant) was most enriched in mutations in the classical HLA-A/B/C genes, B2M, and TAP1/2 (Supplemental Figure 7C). This is likely explained by our findings that ABC tumors – and MCD cell models specifically – display high levels of transcription and translation of APP machinery and are critically dependent upon IRF4 expression, which is activated by MYD88 and BCR signaling (see Figure 5). Thus, the oncogenic signaling required for these tumors leads to high immunogenicity, which is bypassed by inactivating mutations in the major components of the class I pathway.

GCB-classified tumors display high frequencies of mutations in B2M and EZH2. Hyperactive EZH2 was recently correlated with MHC-negative tumors (Ennishi et al., 2019) and is consistent with our data finding EZH2 as a negative regulator via unbiased screens. These tumors also show high levels of IRF8 mutations and B3GNT2 amplifications, mostly stemming from EZB-subtype patients (Supplemental Figure 7C).

Genetic co-occurrences were also characterized (Figure 6E). In some cases, this reflects gene proximity at deleted or amplified chromosomal sites, such as the common loss of CTDSPL2 and PDIA3 in tumors with B2M gene deletions, all located on chromosome 15 (Supplemental Figure 7B). Conversely, some genetic alterations were mutually exclusive – for example, HLA-B mutations were rarely found simultaneously with B2M loss, presumably because one or the other would suffice for immunoevasion (Figure 6E). Loss of the classical MHC class I genes were rarely observed with activating EZH2 mutations, highlighting subtype-specific immunoevasion strategies.

We next asked whether the MHC-I regulators discovered in DLBCL might function similarly in other cancers. Using public sequencing data from The Cancer Genome Atlas (TCGA), we correlated gene expression with tumor immune cell infiltration as measured by CD8+ T cell signatures. Unsurprisingly, each cancer showed a variable pattern of gene/T cell correlation, but ∼30 positive regulator genes showed consistent statistically significant positive correlations with CD8+ infiltration across more than 20 malignancies (green highlighted genes, Figure 6F). Similarly encouraging, ∼10 validated, novel negative MHC-I regulators displayed consistent negative correlation between expression level and T cell signatures (green highlighted genes, Figure 6G). These data demonstrate that genes discovered via unbiased screening in one tumor type to regulate MHC-I are favored to play a similar role in other cancers, thus winnowing potential targets for improving immunotherapies.

### EZH2 and thymidylate synthase are therapeutic targets for DLBCL

Immunotherapy research has largely focused on manipulating tumor infiltrating lymphocytes (TILs), e.g. using checkpoint inhibitors, with relatively less attention on promoting tumor cell immunogenicity by modulating MHC-I regulators. To identify small molecules that enhance surface MHC class I, we conducted a targeted small molecule screen. Compounds were selected by cross referencing validated negative regulators identified by the genetic screens (Figure 3) with small molecule consensus gene targets; we also selected a number of other known immunity-modulating drugs. 48 compounds were tested in 1:4 dilution series on two MHC-I^low^ GCB DLBCL cell lines, DB and SUDHL4. Flow cytometry-based analysis of MHC-I levels 48 hours post-treatment revealed that several compounds targeting EZH2 or thymidylate synthase (TS/TYMS) enhanced surface levels of class I in a dose-dependent manner (Supplemental Figure 8, Supplemental Table 4).

EZH2 is the catalytic core of the PRC2 complex, a general transcriptional repressor involved in methylating K27 of histone H3; small molecule inhibitors (EZH2i) are already being pursued in clinical trials to treat a variety of malignancies (e.g., GSK126 and tazemetostat) (Italiano et al., 2018; Lue and Amengual, 2018). Indeed, inhibiting EZH2 with GSK126 enhanced surface levels of MHC-I in approximately half of 32 DLBCLs tested, whereas MHC-II responses were widely variable and did not trend with MHC-I changes (Figure 7A; Supplemental Figure 9A). As predicted from their already near-maximal levels of MHC-I expression, ABCs were largely unresponsive to EZH2i. Confirming the functional relevance of enhanced MHC-I expression, we expressed NY-ESO-1 in SUDHL4 cells (Figure 7B; Supplemental Figure 9B) and found that either KO of EZH2 (Figure 7C, p < 0.0001) or pre-treatment with the EZH2 inhibitor tazemetostat (Figure 7D, p < 0.0001) enhanced primary CD8+ T cell recognition of the target tumor cells. Mechanistically, we observed that inhibition of EZH2 led to significant loss of repressive H3K27me3 marks at the promoters of NLRC5 and HLA-B in SUDHL4, suggesting that these genes are directly controlled by the EZH2-containing PRC2 complex (Figure 7E, p < 0.0001).

**Figure 7.**
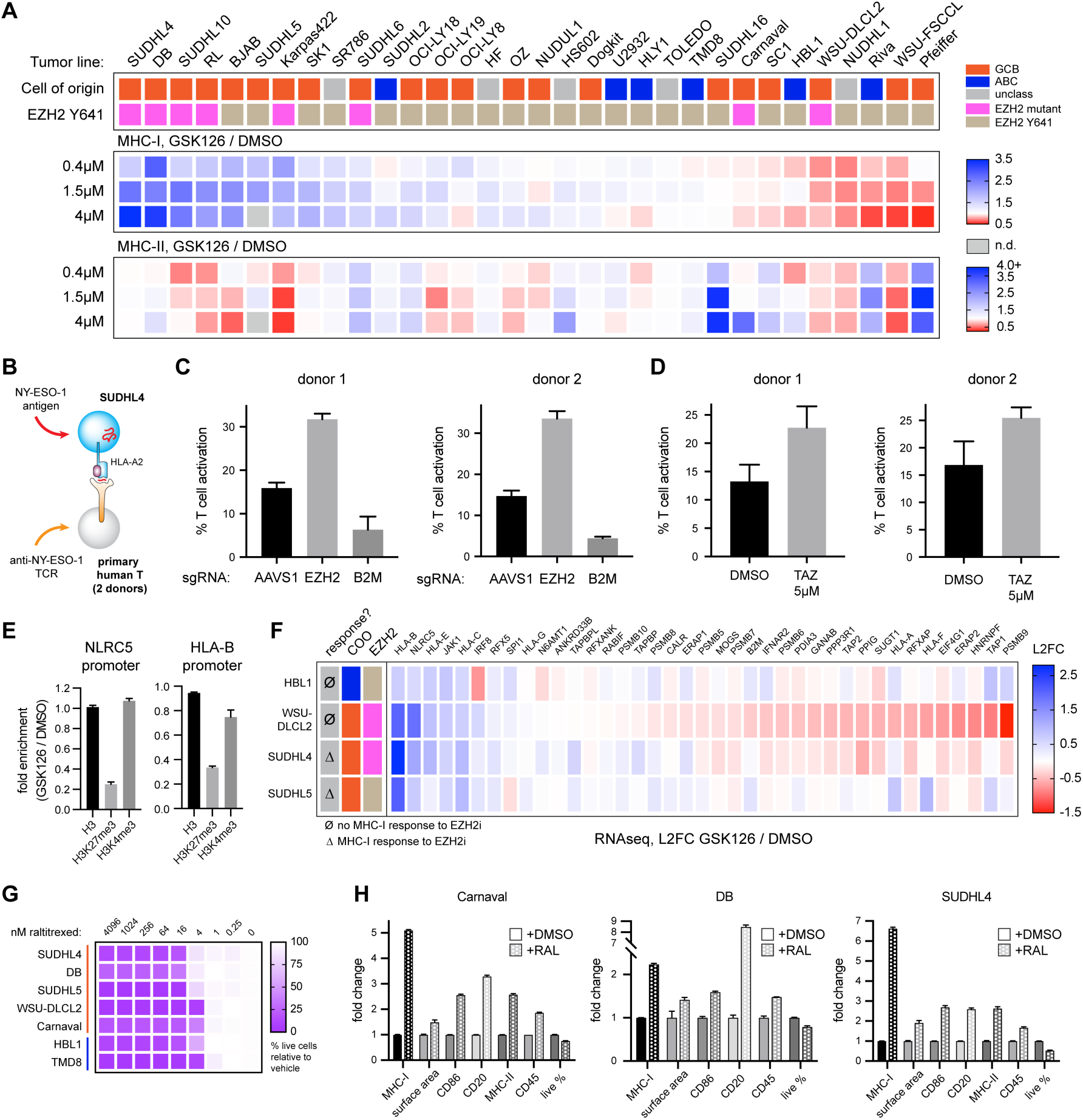
Pharmacological targeting of EZH2 and TS enhances tumor antigen presentation. **(A)** 32 tumor lines were treated with the EZH2 inhibitor GSK126 at the indicated concentrations for 7 days. Average fold increases of MHC-I and MHC-II are plotted by heatmap. At the top, cells are categorized by their cell-of-origin. Targeted sequencing of the EZH2 Y641 region was also conducted to determine mutational status of each line. **(B)** Schematic of T cell activation assay. NY-ESO-1 was transduced into the HLA-A2^+^ SUDHL4 tumor line. Primary human T cells were transduced with a known TCR targeting the NY-ESO-1 peptide 157-165 bound to HLA-A2. T cell activation was measured after co-culture by 4-1BB upregulation. **(C)** SUDHL4-NY-ESO-1 cells were transduced with sgRNAs targeting AAVS1 (safe harbor, negative control), EZH2, or B2M; 9 days later, they were co-cultured with anti-NY-ESO-1 T cells to quantify T cell activation. T cells grown without target cells were set to 0% (9.28% average donor 1; 2.19% average donor 2). **(D)** SUDHL4-NY-ESO-1 cells were treated with either DMSO or 5μM tazemetostat for 5 days, followed by washout of drug and subsequent co-culture with anti-NY-ESO-1 T cells. T cells grown without target cells were set to 0% (5.58% average donor 1, 13.27% average donor 2). **(E)** ChIP for total histone H3, H3K27me3, and H3K4me3 was conducted in SUDHL4 cells with and without GSK126 treatment. Fold of GSK126/DMSO treatment is plotted for the NLRC5 and HLA-B promoter amplicons. **(F)** HBL1, WSU-DLCL2, SUDHL4, and SUDHL5 cell lines were treated with subtoxic doses of GSK126 for 4 days prior to RNA extraction and RNAseq analysis. Plotted are transcript changes with drug treatment of known and newly identified MHC-I regulators. **(G)** The indicated DLBCLs were cultured with serial dilutions of the TS inhibitor raltitrexed to determine % growth inhibition after 48 hours. **(H)** Cells were treated with raltitrexed or vehicle control and stained for various surface markers after 48 hours (Carnaval, 30nM; DB, 20nM; SUDHL4, 75nM). Surface areas were calculated by automated diameter measurements; “% live” indicated relative fraction of cells live by FACS scatter. For entire figure, bar graphs represent mean with standard deviations, minimum n = 3.

How do we account for the variability in GCB tumor responsiveness to EZH2i? As EZH2 mutations at Y641 are enzymatically activating (Yap et al., 2011), this might be predictive of drug sensitivity. Indeed, most of the highly responsive lines are heterozygous at this position (Figure 7A; Supplemental Figures 9C and 9D). EZH2 mutational status is not, however, entirely predictive of class I response, as a number of WT tumors responded well, and two mutant tumors were unresponsive. This mimics results in both preclinical studies (McCabe et al., 2012) and clinical trials (Italiano et al., 2018), where growth inhibition responses to EZH2i were observed in tumors with both mutant and WT EZH2.

Why would a hyperactive EZH2 mutant tumor fail to respond to EZH2 inhibition, especially if PRC2 control of NLRC5 and HLA-B appears to be conserved (Burr et al., 2019; Ennishi et al., 2019; Zingg et al., 2017)? We treated several cell lines with GSK126 and analyzed global transcriptional changes by RNAseq (Figure 7F). Responsive lines displayed increased HLA-B and NLRC5 transcripts, as predicted from ChIP analyses. Interestingly, an MHC-I non-responsive EZH2 mutant tumor, WSU-DLCL2, mirrored these changes but also exhibited downregulation of other genes involved in productive antigen presentation, including a number of the novel regulators identified by our CRISPR screens. Thus, the effect of EZH2 inhibitors on MHC-I expression can be complicated by antagonistic effects on APP accessory genes, and global understanding of regulatory pathways is required to fully appreciate tumor-specific differences.

We also pursued inhibitors of thymidylate synthase, as these showed significant responses in our targeted drug screens. TS contributes to the biosynthesis of thymidine and therefore is critical for DNA replication and repair. Inhibitors of TS such as pemetrexed and raltitrexed have been shown as effective chemotherapeutic agents across a variety of malignancies (Rose et al., 2002). Indeed, we observed sensitivity of DLBCLs to TS inhibition, as cell division and viability significantly suffered within 2 days at low nM doses (Figure 7G; Supplemental Figure 10A). Cell diameter measurement indicated that cell size increases within 2 days, consistent with the G1/S block previously described with TS inhibitors (Berg et al., 2001) (Figure 7H, Supplemental Figure 10B, “surface area”).

In a number of tumor lines, treatment with pemetrexed or raltitrexed also significantly boosted surface levels of MHC-I, an increase which could not be explained by surface area increase alone (Figure 7H, Supplemental Figure 10B). Importantly, TS inhibition could even stimulate increases in surface MHC-I on a tumor line that was resistant to induction by EZH2i (Carnaval), suggesting drug specificity in the ability to manipulate MHC-I.

These results indicate that both EZH2 inhibitors and TS inhibitors may be promising treatments for enhancing DLBCL immunotherapies and further that their clinical efficacy would result from a combination of both direct cytotoxicity with enhanced antigenicity/immunogenicity. Importantly, tumors resistant to one treatment may benefit from other MHC-I-augmenting drugs. More generally, given the heterogeneity of tumors and their pharmacological responses, our work suggests the utility of combination therapy regimens to bypass the diverse immune evasion strategies employed by lymphomas.

## Discussion

Cancer immunotherapy exhibits tantalizing but inconsistent responses across multiple malignancies and patients. Efforts have largely focused on enhancing CD8+ T cell function, but there are also opportunities to improve targetability by increasing tumor MHC-I peptide presentation. Additionally, T cell-mediated therapies are likely to be ineffective at eliminating immunologically invisible tumors. Therefore, better understanding global regulation of APP pathways in diverse cancers is essential for optimizing immunotherapy-based interventions.

Here, we employed CRISPR/Cas9 genome-wide screening to create a global map of MHC-I regulators in DLBCL. Dozens of genes broadly participate in MHC-I biogenesis and turnover. These genes function in diverse pathways including mRNA processing, signaling, trafficking, ER quality control, epigenetics, and translation, among others. Clearly, regulating such a critical immune complex requires an impressive coordination of cellular events.

It is important to consider two caveats of the screens we conducted. First, CRISPR-mediated KO of essential genes will eliminate these cells and their corresponding sgRNAs. For example, we did not identify proteasome subunits as positive regulators of APP, likely because of their essentiality. Second, the phenotype screened – total surface MHC-I – does not report on genes important for the selective presentation of specific peptides. For example, TAPBPR (TAPBPL) is known to play a role in the peptide editing of class I molecules and affects the immunopeptidome repertoire (Hermann et al., 2015), though its loss apparently does not impact overall levels of MHC class I in DLBCL, similar to prior observations in HeLa cells (Boyle et al., 2013).

Although we individually deleted ∼1% of annotated human genes across four tumor lines, our list of validated hits is certain to be incomplete. Determining the relative “importance” of the identified genes in tumor peptide presentation is complicated by multiple factors including sgRNA frameshift efficiency, mRNA and protein half-lives, and genetic redundancies. With these caveats in mind, targeting the well-defined gene products in antigen presentation (e.g. members of the peptide loading complex and RFX family of transcription factors) exerted the strongest negative effects on MHC-I surface expression. We further identify SUGT1 as a strong positive regulator and show that it controls NLRC5, thereby regulating transcription of MHC-I pathway genes. SUGT1 has been shown to interact with ribosome elongation factor eEF1A1 (Novosylna et al., 2015), which binds defective ribosomal products (DRiPs) (Yewdell, 2011) and stimulates their degradation (Gandin et al., 2013; Hotokezaka et al., 2002); therefore, a role for SUGT1 outside MHC transcription (e.g. peptide supply) is certainly possible. How other novel positive regulators influence MHC-I biogenesis is of clear interest and creates opportunities for future research in the APP field.

Relatively less attention has been given to negative regulation of MHC-I in cancer, which potentially occurs at every level of its biogenesis. As tumor MHC-I levels positively correlate with clinical responses (Harel et al., 2019), increasing MHC-I expression provides the obvious, if largely untapped, potential to enhance immunotherapy. Excitingly, we identify dozens of negative regulators in DLBCL that represent potential therapeutic targets. In particular, the large number of clathrin-mediated endocytosis factors and lysosome-directing trafficking proteins strongly suggest mechanisms by which MHC-I is removed from the cell surface in these tumors and warrant investigation in future studies.

Remarkably, we identified a number of genes that co-regulate MHC-I and MHC-II, some in opposing directions. B cells, and ∼75% of DLBCL tumors, constitutively express MHC-II, and nearly all cell lineages can express class II after exposure to IFNγ and other cytokines. CD4+ T cells can participate in tumor clearance by directly lysing tumor cells as well as by enhancing tumor lysis by other immune cell types (Alspach et al., 2019; Sledzinska et al., 2020), and MHC-II may represent an important target for non-conventional CD8+ T cells that can be induced by specific viral vectors (Hansen et al., 2013). As MHC-II may be critically important for DLBCL patient survival (Ennishi et al., 2019), the dual role of some genes we identified is likely clinically relevant. For example, FBXO11 positively regulates MHC-I but negatively regulates MHC-II. In DLBCL, genetic inactivation of FBXO11 stabilizes the proto-oncogene BCL6 (Duan et al., 2012); it would be appropriate to examine class II levels in these cases and whether FBXO11 mutational status could assist in the prediction of clinical responses.

Most gene disruptions in our cellular models of DLBCL affect class I presentation similarly across different tumors, which is somewhat surprising given the heterogeneity across DLBCL (Nissen et al., 2019). However, we did observe genes having variable (and even opposing) effects in different tumors. This could be due to key differences in gene networks, MHC allomorphs, or other polymorphic genes. One striking finding was that ABC-type tumors resisted MHC-I induction compared to GCBs through either genetic manipulation, EZH2 and TS inhibition, or interferon treatment.

Indeed, ABC tumors appear to have constitutively maximal levels of antigen presentation machinery stemming from high mRNA levels. This is partially explained by the ABC tumor transcription factor repertoire, including IRF4, which we show associates with HLA-B promoters, and addiction to NF-κB signaling, which is known to stimulate transcription via the κB1/2 motifs of the MHC-I promoters. We hypothesize that the MHC-I transcription factor NLRC5 also plays an important oncogenic role in ABCs (Ludigs et al., 2015). High expression and signaling of NF-κB and STAT3 is common in ABC tumors (Ding et al., 2008; Lam et al., 2008), and both pathways can drive expression of NLRC5 (Cui et al., 2010; Lu et al., 2018), which we observe is highly expressed in ABCs. Since NLRC5 is generally spared of inactivating mutations in ABC tumors, we speculate that NLRC5’s reported roles in negatively regulating toxic type I interferon responses could be necessary for survival of ABCs and may explain its selective retention even in the face of immune pressure (Cui et al., 2010; Yang et al., 2012).

Our expanded list of MHC-I regulators provides a potential explanation for the frequently identified clinical samples with absent/altered MHC-I levels without accompanying mutations in “canonical” APP genes (Challa-Malladi et al., 2011; Ennishi et al., 2019). Indeed, we now expand the list of targets for immunoediting, many of which may only have subtle effects that may be selected sequentially as alterations combine to gradually increase escape. Gradual evolution may be a favored strategy since tumors face other selection pressures, including NK cells, which can eradicate tumors based on low MHC-I expression.

Additionally, even genes that show minor effects on APP in our single gene KO studies may be much more relevant in the complex genetic environments of tumors (e.g. synergy/addition between MHC allele mutations and trafficking factors). Indeed, most patient samples in our analysis cohort displayed alterations in multiple genes.

ABC patient tumors, which naturally drive class I expression, are unsurprisingly identified as highly immunoedited. Indeed, many ABC model cell lines were unusable in our analyses, as they lack surface MHC-I to study (e.g., OCI-LY10, DLBCL2, OYB). Tumors without such genetic immunoevasion strategies are likely highly immunogenic and must escape from T cell pressure in other ways – their frequent development in extranodal (and often immune privileged) sites is well established (Bruno et al., 2014; Kraan et al., 2013; Shi et al., 2019). Together, these findings might help explain the disappointing clinical outcomes in single-agent immune checkpoint blockade trials for DLBCL (Ansell et al., 2019; Lesokhin et al., 2016) and indicate that NK-mediated or CAR-T immunotherapies may be better suited for ABC DLBCL immunotherapies.

GCB tumors employ different strategies for immunoevasion compared to ABC tumors. Many GCB tumors constitutively express low MHC-I based on epigenetic regulation, as mRNA and protein levels are readily induced by type I and II interferons. Transcription is repressed at least in part by the PRC2 complex, identified recently in a genetic analysis of MHC-I^low^ tumor biopsies (Ennishi et al., 2019). EZH2 Y641 mutations, common in DLBCL, are considered to be hyperactivating (Yap et al., 2011) and are the target for EZH2 inhibitors aimed at starving mutant EZH2 tumors of PRC2 activity in the clinic (Kim and Roberts, 2016; Lue and Amengual, 2018).

Our identification of components of the PRC2 complex as negative class I regulators via unbiased screening supports the role of this complex in immunoevasion. Ennishi et al suggested that the use of EZH2 inhibitors to enhance antigen presentation would be limited to tumors with activating mutations. However, given that our screens were conducted in WT EZH2 GCB lines, even unmutated EZH2 can clearly repress MHC-I. Consistent with these findings, we show that many WT EZH2 GCBs and unclassified tumor lines respond to EZH2 inhibitors with increased MHC-I levels. Conversely, EZH2 inhibitors failed to increase MHC-I in two EZH2 tumors with activating mutations. Together, these findings indicate that first, basal levels of PRC2 activity in WT EZH2 cells can contribute to repression of MHC class I, and second, mutational status of EZH2 does not necessarily predict pharmacological response. This is in line with a recent report of PRC2 regulation in small cell lung cancer and neural progenitors, which do not contain Y641 mutations (Burr et al., 2019).

Indeed, a more holistic understanding of antigen presentation is required to interpret individual tumor responses. As a prime example, we show that while MHC-I transcription in the WSU-DLCL2 tumor line is regulated by its mutant EZH2, productive antigen presentation requires the concerted action of many other regulatory genes. Regardless, clinical trials involving EZH2 inhibition should clearly collect data involving T-cell infiltration, MHC-I/II status and typing, and mutational burden/neoepitope predictions moving forward.

Another therapeutic target identified by our genetic and small molecule screens is TS, important to the biosynthesis of thymidine by catalyzing the methylation of dUMP to dTMP using methylene-THF as a cofactor (Wahba and Friedkin, 1962). Depletion of thymine leads to a cellular response known as thymineless stress and eventual cell death (Cohen, 1971); importantly, TS activity is also crucial during the S phase of cell division, and TS expression is elevated in proliferating cells (Rahman et al., 2004). As its enzymatic function supports DNA replication and repair, TS is a longstanding target for inhibition in anticancer therapies. 5-fluorouracil was introduced in 1957 (Heidelberger et al., 1957), and TS inhibitors are currently approved in treatment of breast, colon and rectum, gastric, gastroesophageal junction, and pancreatic adenocarcinomas as well as non-squamous, non-small cell lung cancers. Still, studies continue to investigate TS catalysis and attempt to improve the functionality of TS inhibitors (Kholodar et al., 2018; Li et al., 2019).

Interestingly, a recent study described enhanced immune control of tumors with combination TS inhibitor (pemetrexed) and anti-PD-L1 blockade, with clear evidence of heightened T cell priming, increased intratumoral leukocytes, and a greater T cell inflamed phenotype (Schaer et al., 2019). Our results are consistent with their observations of tumor immunogenic cell death, and higher levels of MHC class I on transformed cells may provide greater sources of tumor specific antigens. These results provide initial evidence to examine the use of TS inhibitors in DLBCL, perhaps in combination with immune-based therapies.

Though our work was conducted in B cell lymphomas, no doubt some of the newly identified MHC-I regulators will be active in other tumor lineages as well as in normal tissues. These genes could potentially be targeted by viral immunoevasins and may be dysregulated in autoimmunity. We detected significant correlations of gene expression with CD8+ T cell signatures across numerous other human cancers using TCGA data, providing clear avenues to pursue. Sequencing data in other public databases might also be productively mined for clues of the participation of these genes in other diseases involving CD8+ T cell immunosurveillance.

## Acknowledgements

Special thanks to the NIAID Research Technologies Branch, including Tim Myers, the Genomic Technologies facility, Teresa Hawley, and the Flow Cytometry facility. Thanks to the NCI Center for Cancer Research sequencing facility and staff, as well as support personnel in the laboratories of L.M.S and J.W.Y. We also acknowledge the Centre for Clinical Immunology and Biomedical Studies at the Institute for Immunology & Infectious Diseases for HLA typing. This research was supported by the Intramural Research Program of the NIH – NCATS, NIAID, and NCI – and specifically the NCI FLEX award to J.W.Y and L.M.S.

## Author Contributions

Conceptualization, D.D., J.D.P, L.M.S., and J.W.Y.; Methodology, D.D., J.D.P., L.M.S., J.W.Y.; Formal Analysis, T.E.M., N.F., G.W.W., D.W.H.; Investigation, D.D., J.D.P., B.W., M.E.G., J.H.A., J.H., R.J.K., T.E.M., M.O.S.; Resources, R.J.K., M.C., C.J.T., N.P.R.; Data Curation, T.E.M., N.F., G.W.W., D.W.H.; Writing – Original Draft, D.D., J.W.Y.; Writing – Review & Editing, J.D.P., M.E.G., J.H.A., J.H., T.E.M.; Visualization, D.D., J.D.P., M.E.G.; Supervision, C.J.T., J.B.L., N.P.R., T.M.K., L.M.S., J.W.Y.

## Methods

### DLBCL cell culture

All DLBCL cell lines were cultured at 37°C in 5% CO_2_-containing humidified incubators using Advanced RPMI (Gibco) containing 5% fetal bovine serum (Hyclone or Seradigm, heat inactivated and tet-tested), 1% penicillin/streptomycin (Gibco), and Glutamax (Gibco). Lines were routinely tested for mycoplasma using either the Mycoalert Mycoplasma Detection Kit (Lonza) or the Universal Mycoplasma Detection Kit (ATCC). Cell identity was confirmed by either STR testing (ATCC) or copy number variant fingerprinting of 16 loci from cell-derived genomic DNA (Jonathan Keats, personal communication). The generation of doxycycline-inducible Cas9 clones is described in (Phelan et al., 2018).

### sgRNA cloning

pLKO.1-puro (Addgene #52628, a gift from Scot Wolfe) and the modified pLKO.1-puro/GFP vector system described in (Phelan et al., 2018) were used to deliver individual sgRNAs to Cas9-expressing cells. Empty vector was prepared by digestion with BfuAI (New England Biolabs) at 50°C for 3 hours and heat inactivation at 65°C for 25 minutes. This was followed by dephosphorylation via Antarctic Phosphatase (New England Biolabs) for 1.5 hours at 37°C and heat inactivation for 4 minutes at 80°C. Digested, dephosphorylated vector was gel purified according to manufacturer’s protocol (Qiagen Gel Extraction kit). Overhang DNA oligos corresponding to sgRNA sequences (ACCG flank on 5’ end, and CAAA flank on the reverse complement 3’ end) were synthesized by Eurofins and mixed equimolar at a final concentration of 100μM. Oligos were phosphorylated by T4 PNK (New England Biolabs), annealed by temperature drop, diluted, and ligated into pLKO.1 vectors using T4 DNA ligase and manufacturer protocols (New England Biolabs). DNA was transformed into Stbl3 bacteria for plasmid preparation (Thermo Fisher).

### sgRNA library

The Brunello CRISPR library targeting the human genome was obtained from Addgene (via John Doench and David Root, Addgene #73178), and 400ng was electroporated into Stbl4 bacteria (Thermo Fisher). Colonies were grown by incubation at 30°C on large bioassay plates; bacteria were harvested by scraping colonies in cold LB medium. DNA was purified using HiSpeed Maxi prep kits (Qiagen).

### Lentivirus/retrovirus production

293FT were used to prepare all lentiviruses and retroviruses and were maintained at 37°C in 9% CO_2_-containing humidified incubators using DMEM (Gibco) containing 10% fetal bovine serum (Hyclone, not heat inactivated) and non-essential amino acids (Gibco). Cells were plated one day prior such that transfection was conducted at a confluency of 70-90%. For Brunello library virus preparation, library DNA, pMD2.G (Addgene #12259, gift from Didier Trono), and psPAX2 (Addgene #12260, gift from Didier Trono) were transfected at a 4:3:1 ratio using TransIT293 reagent into T225 flasks as indicated by the manufacturer (Mirus Bio). For individual sgRNA preparations, 60mm plates were transfected with pLKO.1 plasmid, pMD2.G, and psPAX2 at a ratio of 2:1.5:1.06 using TransIT293. Virus-containing supernatant was collected at 24 and 48 hours post-transfection, spun at 500g for 10 minutes, and incubated with Lenti-X concentrator (Takara) for at least 24 hours. Virus was concentrated 32x, resuspended in PBS, aliquoted, and frozen at - 80°C. Brunello virus was titrated by puromycin selection compared to uninfected controls; cell viability was measured by flow cytometry as a function of virus used. Production of the MuLV retroviruses was similar to that of lentiviruses, using MSCV-IRES-GFP (Addgene #20672, gift from Tannishtha Reya) as an expression backbone. MSCV vector containing either nothing (empty vector) or the CTAG1A gene were co-transfected in 293FT with pMD2.G and pUMVC (Addgene #8449, gift from Bob Weinberg) at a ratio of ∼1.5:1.2:1.8. Retrovirus was collected, purified, and concentrated identically to lentiviruses.

### Genome-wide screening for MHC-I

Two replicates of each dox-inducible Cas9 cell line were independently transduced with Brunello library lentivirus such that transduction efficiency was between 15-25%. sgRNA coverage was maintained throughout the experiments at >500 copies of each sgRNA (∼40 million cells for the ∼77,000 sgRNAs). Puromycin was added after 3 days of infection, and resistant cells were grown out for 8 days. Pre-doxycycline input samples were harvested, and 200ng/mL doxycycline was added for 8-11 days to induce Cas9 and initiate genetic ablations. Cells were passaged every two days with fresh medium containing doxycycline until the first round of sorting, at which point a post-doxycycline input sample was harvested. 65 million cells were stained with the anti-pan-MHC-I antibody W6/32 (purified antibody from BioXcell, directly conjugated with AlexaFluor 647 via Molecular Probes kit), washed 3x in RPMI/FBS-containing staining buffer, and sorted for the lowest and highest ∼5% populations. Sorted cells were placed back into culture in conditioned medium and expanded continually for 1 week prior to resorting. Resulting cells were stained and sorted for the lowest of the low and highest of the high (∼15%) MHC-I expressors, yielding ∼2-4 million cells for each final population. gDNA was extracted from all input and sorted samples using either DNeasy kits (Qiagen) or QIAmp DNA Blood maxi kit (Qiagen).

### CRISPR library preparation and sequencing

Sequencing libraries were prepared as previously described (Phelan et al., 2018; Webster et al., 2019). Briefly, sgRNA sequences were amplified from 240μg of genomic DNA per sample with primers 5’AATGGACTATCATATGCTTACCGTAACTTGAAAGTATTTCG and 5’GTAATTCTTTAGTTTGTATGTCTGTTGCTAT TATG and ExTaq (Takara) polymerase. In total, 24 PCR replicates were performed per sample in 100μL reactions with 10μg of genomic DNA per replicate. FACS-sorted populations contained fewer cells and thus all recovered genomic DNA (5-40μg) was amplified in 100μL reactions. Primary PCR reactions were pooled, and sequencing adapters and sample indices were added to 5μL of primary PCR product. Reactions were amplified for 24-27 cycles and then size selected using 2% E-Gel EX Size Select gels (Invitrogen). Libraries were quality checked using Agilent 2100 bioanalyzer (Agilent), Kapa qPCR (Kapa Biosystems), and quantitated by QuBit high sensitivity standards (Thermo Fisher Scientific). Libraries were sequenced on an Illumina NextSeq500 achieving an average of >340X coverage in unsorted samples and >80X coverage in flow-sorted samples. Custom perl scripts were used to extract sgRNA sequences from fastq files and bowtie2 was used to align sgRNA sequences allowing for a 1bp mismatch (Phelan et al., 2018).

### STARS analysis of screen

To assemble sgRNAs and rank genes from the screening data, we used STARS_v1.2 (Doench et al., 2016). First, we added 250 pseudo “genes” based on four randomized non-targeting guides present in the Brunello library. Additionally, the most poorly sequenced/expressed sgRNAs in the input samples were removed from analysis. Z-scores for each sgRNA were calculated based on Log_2_FoldChange of sorted populations to input controls. Segregation scores were generated by the difference in Z-score of a sgRNA in MHC-I high vs MHC-I low. For robustness, STARS score calculation excluded each gene’s first ranking perturbation, and the perturbation percentage threshold was set to 25%, for an increased number of valid gene guides (up to 4 per gene). Guide ranking was based on their duplicate-average segregation score. STARS analysis was carried out on python 2.7. Note that STAR p-values rounded to 0 were manually set at -log(p-value) of 8. We combined all hits showing a p-value of <0.01 or a segregation score outside the range of -1.5 – 1.5 (Supplemental Table 2). For all STRING analyses, confidence was set to “medium”, with line thickness indicating relative confidence in the genetic interaction (Szklarczyk et al., 2019).

### Validation of MHC-I regulators and flow cytometry

DLBCLs were plated in flat-bottom 96-well plates and infected with saturating amounts of concentrated sgRNA-expressing lentiviruses. Non-targeting sgRNAs were delivered to wells of cells separately from targeting sgRNAs (NT or AAVS1 in pLKO.1-puro if the experimental sgRNAs were in pLKO.1-puro/GFP and vice versa). Two days post-infection, cells were split into medium containing 400ng/mL doxycycline and puromycin (0.5-2 μg/mL depending on the line). Cells were passaged in constant doxycycline and puromycin for ∼7-9 days (9-11 days total post-infection) prior to analysis. Cells were harvested and NT sgRNA infected cells were mixed with experimental KO lines as internal controls delineated by GFP expression. Cells were washed into lymphoma staining buffer (RPMI without phenol red supplemented with Glutamax and 1% FBS) and subsequently stained with antibody solutions at 4°C for 30 minutes with slight shaking. Cells were washed 2-3x prior to analysis on a Celesta, Fortessa X-20, or FACSCalibur (BD Biosciences). FCS files were analyzed by Flowjo version 10 (BD Biosciences). Changes in MHC-I surface expression were calculated as either the changes in raw MFI, or the derived parameter of MHC-I per CD147 per cell. All samples were normalized to NT sgRNA-infected cells. Antibodies used throughout this work include the anti-pan-MHC-I antibody W6/32 (purified antibody from BioXcell, directly conjugated with AlexaFluor 647 via Molecular Probes kit); anti-CD147 (clone HIM6, BV421, BD); anti-HLA-DR (clone L243, PE-Cy7, eBioscience); anti-HLA-DR,DP,DQ (clone Tu39, FITC, BD); anti-HLA-A2 (clone BB7.2, APC, BD); anti-IFNGR (clone GIR-94, BD Pharmingen).

### Statistical analyses

One-way ANOVA or unpaired t tests were performed using GraphPad Prism 8.3 (GraphPad Software LLC). One-way ANOVA tests were adjusted for multiple comparisons by Dunnett’s test. Statistical tests and p-values are summarized in Supplemental Table 5.

### Western blotting and immunoprecipitation

When indicated, cells were treated with 500 U/mL human IFNγ and human IFNβ for 2 days prior to isolation (Peprotech). For all lysate blots, cells were harvested and washed in PBS prior to direct lysis with SDS lysis buffer at 95°C for 10-15 minutes - 50mM Tris pH7.4, 150mM NaCl, 1mM EDTA, 2% SDS, protease inhibitor cocktail (Roche), and 15U/mL DNase I (New England Biolabs). Protein concentration of all lysates was determined by DC Protein Assay (Bio-Rad), absorbance measured by a Synergy H1 plate reader (BioTek). For samples subjected to immunoprecipitation, cells were washed in PBS and lysed in RIPA buffer (25mM Tris pH7.4, 150mM NaCl, 1% NP-40, 0.5% sodium deoxycholate, 0.1% SDS, protease inhibitor cocktail) with gentle mixing for 15 minutes at 4°C. Debris was pelleted by centrifugation at 15,000g for 15 minutes. 800-900μg protein was typically used per IP with an anti-CIITA antibody (Cell Signaling). After overnight incubation, magnetic protein G particles (ThermoFisher) were used to isolate captured CIITA. Both lysate and IP samples were prepared in 4X NuPAGE LDS sample buffer (Invitrogen) and run on 4-12% NuPAGE Bis-Tris mini or midi gels. Transfer to nitrocellulose was conducted using iBlot or iBlot 2 (Life Technologies) as described by manufacturer. Blots were blocked using Odyssey Blocking Buffer (OBB, Licor) and probed overnight in primary antibodies in OBB with 0.1% Tween-20. Secondary antibodies were incubated for 1 hour at room temperature in OBB with 0.1% Tween-20. Blots were scanned using the Licor Odyssey CLx (Licor) and analyzed using ImageStudio (Licor). Antibodies used for Western blotting include: anti-SUGT1 (Abcam); anti-NLRC5 (clone 3H8, EMD Millipore); anti-HSP90 (Santa Cruz); anti-HLA-A/B/C (clone EMR8-5, Abcam); anti-histone H3 (Cell Signaling); anti-GAPDH (Proteintech); anti-MHC-II (clone LGII-612.14, Cell Signaling); anti-CIITA (clone 7-1H, Santa Cruz); anti-B2M (Dako); anti-TAP1 (EMD Millipore); anti-IRF4 (Cell Signaling); anti-phospho-JAK2 (Cell Signaling); anti-14-3-3 (Santa Cruz).

### HLA typing

HLA typing for class I (HLA-A/B/C) and class II (DQA1; DQB1, DRB1 3,4,5; DPB1) was performed by an American Society for Histocompatibility and Immunogenetics (ASHI)-accredited laboratory at The Institute for Immunology and Infectious Diseases at Murdoch University Western Australia using locus-specific PCR amplification of genomic DNA. The assay has been adapted from previously published protocol for Barcoded-PCR method (Erlich et al., 2011) with modifications to the primer sequences. Briefly, 11 PCR amplifications per sample targeting the different HLA loci were set up with primers for a given sample tailed with a specific barcode tag sequence. Amplified products were quantitated, normalized, and pooled by subject up to 48 subjects. The pooled and normalized PCR reactions were purified using 1.8x the PCR reaction volume of AMPure XP beads (Beckman Coulter Inc). Samples were prepared for sequencing on Illumina MiSeq using the manufacturer’s standard library preparation protocol. These libraries were quantified using Kapa universal qPCR library quantification kits (Kapa Biosystems). Sequencing was performed on an Illumina MiSeq using the 2 × 300 paired-end chemistry kit (Illumina). Reads were quality-filtered and passed through a proprietary allele calling algorithm and analysis pipeline using the latest IMGT HLA allele database (Robinson et al., 2015) (http://www.ebi.ac.uk/ipd/imgt/hla/) as a reference.

### RNAseq

RNA was purified by either TRIzol extraction (Invitrogen) or RNeasy kits (Qiagen) according to manufacturers. All samples had an RNA Integrity (RIN) score of >9 as determined by the Agilent 2100 bioanalyzer or Agilent 2200 Tapestation system (Agilent). For RNAseq of cell lines (Figure 5C), polyA selected mRNA libraries were generated using the Illumina TruSeq Stranded Library protocol as described by the manufacturer. Briefly, 100ng to 1μg of total RNA was used as the input to an mRNA capture with oligo-dT coated magnetic beads. The mRNA was fragmented, and then random-primed cDNA synthesis was performed. The resulting double-strand cDNA was used as the input to a standard Illumina library prep with end-repair, adapter ligation and PCR amplification. mRNA samples were pooled and sequenced as paired-end 76 base pair reads on a Nextseq 500 running RTA 1.18.64 software. Demultiplexing was done using bcl2fastq v2.17. Both reads of each sample were trimmed for contaminating adapters and low-quality bases using Trimmomatic v0.36 and aligned to the Human hg38 reference genome and Gencode v30 annotation using STAR v2.6.1c. RSEM v1.3.0 was used for gene-level expression quantification. Read- and alignment-level quality was assessed using MultiQC v1.7 (http://multiqc.info/) to aggregate QC metrics from FastQC (http://www.bioinformatics.babraham.ac.uk/projects/fastqc/), FastQ Screen (https://www.bioinformatics.babraham.ac.uk/projects/fastq_screen/), Picard v2.4.1, RSeQC (http://rseqc.sourceforge.net/) and Trimmomatic. For RNAseq of GSK126 treated cells (Figure 7F), libraries were prepared with New England Biolabs product NEBNext Poly(A) mRNA Magnetic Isolation Module, New England Biolabs NEBNext Ultra II Directional RNA Library Prep Kit for Illumina, and NEBNext Multiplex Oligos for Illumina (Dual Index Primers Set 1) using 25ng total RNA input per sample. Library validation was performed on the Agilent 2200 Tapestation System (Agilent) to verify library size and purity. Library quantification via qPCR was performed on the Applied Biosystems 7900HT Fast Real-Time PCR System (ThermoFischer Scientific) using KAPA Biosystems Complete Library Quantification Kit for Illumina. PhiX was added at 1% to serve as an internal control. The resultant final library pool was 1.8pM final concentration with 1% PhiX spike-in. Paired-end sequencing was completed on an Illumina NextSeq 500 system, running Illumina NextSeq Control Software System Suite version 2.2.0 and RTA version 2.4.11. The final library pool was sequenced via 2 x 76 bp run configuration using the NextSeq 500/550 High Output v2.5 kit, 75. Demultiplexed reads were trimmed, aligned, and mapped to reference genome version hg38 using CLC Genomics Workbench software version 12.0. Log2 of total exon counts or TPM were normalized by upper quartile scaling and ANOVA statistics computed using JMP/Genomics version 9.1.

### ChIP

#### IRF4 ChIP

Lysates of crosslinked cells were sonicated in lysis buffer (20 mM Tris-HCl pH8.1, 10 mM EDTA, 0.2% SDS, complete protease inhibitor cocktail, 1 mM NaF, 0.1 mM Na_3_VO_4_, and 10 mM glycerophosphate) using a Covaris Sonicator (8 m, 10% duty cycle, 200 burst per cycle, 75 W peak incident power) to obtain DNA fragment averaging 500-700 bp. Soluble chromatin fraction was diluted 1:2.5 (20 mM Tris-HCl pH8.1, 2 mM EDTA, 250 mM NaCl, 1.6% Triton X-100, complete protease inhibitor cocktail, 1 mM NaF, 0.1 mM Na_3_VO_4_, and 10 mM glycerophosphate) and immunoprecipitation was conducted using 5 µg antibody [IRF4 (Abcam ab101168) or control IgG (Sigma-Aldrich 12-370)] and 1-1.4 mg of chromatin at 4°C for 16 h. Immunoprecipitates were incubated with 25 µl of Dynabeads Protein G (Invitrogen) for 2 h, washed 2x with Low-salt wash buffer (20 mM Tris-HCl pH8.1, 2 mM EDTA, 150 mM NaCl, 0.08% SDS, 1% Triton X-100), 1x in High-salt buffer (20 mM Tris-HCl pH8.1, 2 mM EDTA, 500 mM NaCl, 0.08% SDS, 1% Triton X-100), 1x in LiCl buffer (10 mM Tris-HCl pH8.1, 1 mM EDTA, 1% NP40, 250 mM LiCl), and 1x in TE (10 mM Tris-HCl pH8.1, 1.2 mM EDTA). Chromatin was eluted for two cycles of 30 m at 65°C (1% SDS, 0.1 M NaHCO3), NaCl added to 0.25 M, and heated for 12 h at 65°C to reverse crosslinks. Eluants were treated with Proteinase K and RNaseA, and the DNA was purified using ChIP DNA Clean & Concentrator Kit (Zymo Research). Recovered DNA was analyzed in triplicate by qPCR.

HLA-B tss (5-CGTCACGAGTATCCTGGAAGAA-3, 5-AGGGTCTCAGGCTCCGA-3)

TLR4 tss (5-AATCACCGTCATCCTAGAGAGTTACAA-3, 5-TCTGACCTCTGCCTGGGCTTGGTGAAT-3) (Yang et al., 2012)

SUB1 tss (5-CTTAGAGAACCGAAACCCAAACCTACA-3, 5-TGCAACCCTTCCTGCTTTAACAAGTTT-3) (Yang et al., 2012)

#### H3, H3K27me, and H3K4me3 ChIP

Lysates of crosslinked cells were sonicated in lysis buffer (10 mM Tris-HCl pH8.1, 150 mM NaCl, 5 mM EDTA, 0.5% Sarkosyl, 0.1% sodium deoxycholate, complete protease inhibitor cocktail, 1 mM NaF, 0.1 mM Na_3_VO_4_, and 10 mM glycerophosphate) using a Bioruptor Plus (12 cycles of 30s on/off, high power) to obtain DNA fragment averaging 500-700 bp. Soluble chromatin fraction was diluted 1:5 (10 mM Tris-HCl pH8.1, 5 mM EDTA, 150 mM NaCl, 1% Triton X-100, complete protease inhibitor cocktail, 1 mM NaF, 0.1 mM Na_3_VO_4_, and 10 mM glycerophosphate) with the addition of 40 ng Spike-in Chromatin per reaction (Active Motif 53083). Immunoprecipitation was conducted using 4 µg of anti-H3 (Abcam ab1791), 4 µg anti-H3K27me3 (Sigma-Aldrich 07-449), 4 µg anti-H3K4me3 (Sigma-Aldrich 07-473), 4 µg control IgG (Sigma-Aldrich 12-370), 2 µg Spike-in Antibody (AB_2737370) and 900 µg of chromatin at 4°C for 16 h. Immunoprecipitates were incubated with 35 µl of Dynabeads Protein G for 2 h, washed 2x with Low-salt wash buffer (20 mM Tris-HCl pH 8.1, 2 mM EDTA, 200 mM NaCl, 1% Triton X-100), 2x in LiCl buffer (10 mM Tris-HCl pH8.1, 2 mM EDTA, 1% NP40, 250 mM LiCl), and 1x in TE. Chromatin was eluted and purified as in the IRF4 ChIP experiments.

NLRC5 promoter (5-GGTGATGCCTGCAGAAGTAT-3, 5-CAGCGTTCGCTCCTATTCA-3)

HLA-B promoter (5-TGTTTCTCTGTTCCTCTTGTCC-3, 5-TTGAAGGACATCTATGCTGGATATAG-3)

### qPCR

RNA was harvested from cells using RNeasy kits (Qiagen), including the optional DNase I treatment step. RNA concentration was quantified, and 100ng was used for cDNA synthesis by AccuScript High Fidelity 1^st^ Strand cDNA synthesis kit (Agilent) using the manufacturer’s protocol. qPCR was conducted in triplicate in a total of 20μL using PowerUp 2x SYBR Green master mix (Applied Biosystems), with a 60°C extension temperature, using the Quant Studio 3 instrument (Applied Biosystems). Data was analyzed by standard ΔΔCt method.

Pan-HLA-A (for 5-GCTCCCACTCCATGAGGTAT-3; rev 5-AGTCTGTGACTGGGCCTTCA-3) (Ramsuran et al., 2015)

Pan-HLA-B (for 5-ACTGAGCTTGTGGAGACCAGA-3; rev 5-GCAGCCCCTCATGCTGT-3) (Ramsuran et al., 2015)

Actin (for 5-CATGTACGTTGCTATCCAGGC-3; rev 5-CTCCTTAATGTCACGCACGAT-3)

HLA-DRA (for 5-GGGTCTGGTGGGCATCATTA-3; rev 5-CCATCACCTCCATGTGCCTT-3)

NLRC5 (for 5-CTTTCAGTTTCGTGCAGAGCG-3; rev 5-AGCCAGCCTTGGTCTCCT-3)

### Genetic analysis of DLBCL patient cohort

All primary genetic data is available through the NIH dbGAP system under accession numbers phs001444, phs001184 and phs000178 (https://www.ncbi.nlm.nih.gov/projects/gap/cgi-bin/study.cgi?study_id=phs001444.v1.p1). Mutation calls, DNA copy number analysis and gene expression values were generated as previously described (Schmitz et al., 2018). Somatic mutations were included that displayed a mutant allele frequency greater than 10% and were not found in internal laboratory control DNA or in dbSNP (version 138). Mutations per patient and statistical analyses for mutual exclusivity or co-occurrence were calculated using oncoprinter (https://www.cbioportal.org/oncoprinter) (Cerami et al., 2012; Gao et al., 2013).

### T-cell signature analysis in pan-cancer

Raw data for cohorts with >50 patients were downloaded from The Cancer Genome Atlas (Hoadley et al., 2018). CD8+ T-cell enrichment scores for each tumor were identified by Xcell (Aran et al., 2017). TPM scores for each gene were calculated by cBioportal (cgdsr_1.2.10) (Cerami et al., 2012; Gao et al., 2013). For each gene, cohorts were split into tertiles based upon TPM scores. CD8+ T-cell enrichment score was compared between tumors with the highest expression and those with the lowest expression for direction of change and statistical difference using a two-tailed Wilcoxon Rank Sum Test. P-values were adjusted by Bonferroni correction to correct for multiple analyses within the same cohort.

### T-cell assays

Primary anti-NY-ESO-1^157-165^ human T-cells were generated as described (Patel et al., 2017). Briefly, primary lymphocytes were stimulated with IL-2 and anti-CD3 and retrovirally transduced with the anti-NY-ESO-1 TCR from clinical grade retroviral supernatants via RetroNectin (Takara Bio). Cells were expanded by a rapid expansion protocol involving soluble OKT3, IL-2, and irradiated feeder cells. T-cells were maintained at 37°C in 5% CO_2_-containing humidified incubators in RPMI containing 10% FBS (Hyclone), sodium pyruvate (Sigma), non-essential amino acids (Gibco), Glutamax (Gibco), penicillin/streptomycin (Gibco), and 300 IU/mL IL-2. For SUDHL4, which did not express endogenous NY-ESO-1, cells were transduced with MSCV-IRES-GFP retroviruses containing either NY-ESO-1 or no insert. One week post-infection, stable GFP+ cells were sorted and expanded. Co-culture experiments were conducted in round-bottom 96-well plates with 30,000 or 45,000 effector T-cells and variable numbers of target DLBCL cells. Target cells were typically labeled with Cell Trace Violet (Invitrogen) for 15 minutes in DPBS at 37°C and washed prior to co-culture. Cells were incubated overnight for 12-15 hours and spun into antibody solutions in staining buffer (RPMI without phenol red supplemented with Glutamax and 1% FBS) at 4°C for 30 minutes with slight shaking. Cells were washed 2x prior to analysis by a Fortessa X-20 (BD). Dead cells were gated out by inclusion of SYTOX Blue or SYTOX AAdvanced dyes (Invitrogen) during the last wash step. DLBCL target cells were detected by either GFP expression (pLKO.1-puro/GFP transduction), staining with anti-HLA-DR,DP,DQ (clone Tu39, FITC, BD), Cell Trace Violet fluorescence, or a combination thereof. T-cells were identified and analyzed by staining for CD3 (clone OKT3, BV785, BioLegend); mTCRβ to detect the population of transduced cells expressing the chimeric mouse/human anti-NY-ESO-1 TCR (clone H57-597, PE, eBioscience); and 4-1BB/CD137 to monitor T-cell activation (clone 4B4-1, APC, BD).

### Targeted small molecule screen

The 48 selected small molecules were dry-spotted into ultra-low attachment 384-well plates (Corning) by acoustic dispensing with an Echo 555 Liquid Handler (Labcyte Inc). Briefly, each compound was dry-spotted in a 7-doses, 1:4 dilution series, corresponding to 5.7nM – 23.5 μM final concentrations post cell addition. Each 384-well plate also included columns of 200nL DMSO controls and two empty columns for manually addition of positive and negative controls. Growth tests were first performed for DB and SUDHL4 cells to determine optimal growth conditions in 384-well plates. Cells were then plated in drug-spotted plates in a total volume of 85μL with 33% conditioned medium. For positive controls, 500 U/mL IFNγ (Peprotech) was added to one column of wells one day prior to collection. 5 μg/mL brefeldin A (Biolegend) was added to one column of wells 18 hours prior to collection to serve as negative controls. After a total of 48 hours of drug exposure, cells were stained with anti-pan-MHC-I antibody W6/32-AlexaFluor647, washed 2x with lymphoma staining buffer, and analyzed by a High Throughput Sampler-equipped Fortessa (BD). To calculate fold changes, MHC-I levels were normalized to the average of 16 DMSO control wells.

### Targeted sequencing of EZH2

Genomic DNA was purified from each DLBCL cell line using DNeasy Blood & Tissue Kit (Qiagen) following manufacturer instructions. The genomic region containing EZH2 Y641 (chromosome 7 genomic coordinates: 148,811,063-148,812,103) was amplified by PCR with the primer pair: Fwd: 5’-TGGTAAAGCTCTTGTTCTCCC-3’, Rev: 5’-AGAGTGATTTGGTGGTGTCC-3’; amplicon size: 1041 bp. 50-100 ng of gDNA was amplified in 50 µl reaction using Q5® Hot Start High-Fidelity 2X Master Mix (New England BioLabs) with initial denaturation at 98°C for 30 sec followed by 35 cycles of 98°C 10 sec, 64°C 30 sec, 72°C 40 sec and final extension at 72°C for 2 min. PCR products were purified with DNA Clean & Concentrator-25 kit (Zymo Research) and 5 μl of purified products were loaded on 2% E-Gel EX (Invitrogen) to verify the reaction performance and amplicon size. The rest of the purified PCR product was sent for Sanger sequencing (Eurofins Genomics) with sequencing primer: 5’-CAGCTTTCACGTTGACTG-3’. Chromatograms were manually evaluated using Geneious Prime software.

### EZH2 and TS inhibition assays

DLBCL cell lines plated in 96-well plates were treated with the indicated concentrations of GSK126 (Cayman Chemical), tazemetostat (EPZ-6438, Cayman Chemical), or DMSO vehicle (ATCC) for a total 7 days, splitting every 2-4 days as required with fresh inhibitors. The indicated concentrations of inhibitors were used as they are generally subtoxic doses for most lines. Cells were stained as described above and analyzed by a Fortessa X-20 or FACSCalibur and FlowJo (BD). The fold change compared to DMSO control is an average of 3-4 biological replicates conducted on different days. For TS inhibition assays, pemetrexed (Abcam) or raltitrexed (ApexBio) were used at indicated concentrations for only 2 days without change of medium prior to flow cytometry analysis. For cell growth inhibition assays, Sytox Blue was used to distinguish dead cells and live cell numbers were normalized to CountBright Absolute Counting Beads spiked into the cultures (ThermoFisher).

