## Supplemental figures for "Genome-wide Screens Identify Lineage- and Tumor Specific-Genes Modulating MHC-I and MHC-II Immunosurveillance in Human Lymphomas"

| Cell line | Collection Date | HLA-A Condensed/Putative Allele Result | HLA-B Condensed/Putative Allele Result | HLA-C Condensed/Putative Allele Result | HLA-DPB1 Condensed/Putative Allele Result | HLA-DQA1 Condensed/Putative Allele Result | HLA-DQB1 Condensed/Putative Allele Result | HLA-DRB1 Condensed/Putative Allele Result | HLA-DRB3 Condensed/Putative Allele Result | HLA-DRB4 Condensed/Putative Allele Result | HLA-DRB5 Condensed/Putative Allele Result |
|---|---|---|---|---|---|---|---|---|---|---|---|
| BJAB | 6/12/2018 | 01:01:01G + 02:01:01G | 13:02:01G + 35:01:01G | 04:01:01G + 06:02:01G | 04:02:01G + 04:02:01G | 01:02:01G + 05:01:01G | 03:01:01i + 06:04:01i | 12:01:01i + 13:02:01i | 02:02:01i + 03:01:01i | NP + NP | NP + NP |
| DOHH2 | 6/12/2018 | 01:01:01G + 01:01:01G | 08:01:01G + 44:02:01G | 07:01:01G + 07:04:01G | 02:01:02G + 03:01:01G | 01:01:01G + 01:02:01G | 05:01:01i + 06:02:01i | 01:01:01i + 15:01:01i | NP + NP | NP + NP | 01:01:01i + NP |
| HBL1 | 6/12/2018 | 02:06:01G + 02:06:01G | 51:01:01G + 51:01:01G | 14:02:01G + 14:02:01G | 04:02:01G + 04:02:01G | 01:01:01G + 01:01:01G | 05:03:01i + 05:03:01i | 14:01:01i + 14:01:01i | 02:02:01i + 02:02:01i | NP + NP | NP + NP |
| HLY1 | 28/01/2019 | 02:01:01G + 03:01:01G | 40:01:01G + 44:02:01G | 03:04:01G + 16:04:01G | 04:01:01G + 10:01:01G | 04:01:01G + 05:01:01G | 03:01:01i + 04:02:01i | 08:01:01i + 11:04:01i | 02:02:01i + NP | NP + NP | NP + NP |
| Karpas422 | 8/03/2019 | 02:01:01G + 03:01:01G | 07:02:01G + 27:05:02G | * 07:02:01G + 07:02:01G | 02:01:02G + 04:01:01G | 01:02:01G + 03:01:01G | 03:01:01i + 05:02:01i | 04:01:01i + 16:01:01i | NP + NP | 01:01:01i + NP | 02:02:01i + NP |
| LY19 | 28/01/2019 | 02:01:01G + 02:01:01G | 15:01:01G + 15:01:01G | 03:03:01G + 03:03:01G | 04:02:01G + 04:02:01G | 03:01:01G + 03:01:01G | 03:02:01i + 03:02:01i | 04:01:01i + 04:01:01i | NP + NP | 01:01:01i + 01:01:01i | NP + NP |
| MC116 | 28/01/2019 | 01:01:01G + 03:01:01G | 08:01:01G + 35:01:01G | 04:01:01G + 07:01:01G | 04:01:01G + 04:02:01G | 01:03:01G + 03:01:01G | 03:01:01i + 06:03:01i | 04:01:01i + 13:01:01i | 02:02:01i + NP | 01:01:01i + NP | NP + NP |
| OCILY1 | 6/12/2018 | 02:01:01G + 02:01:01G | 44:02:01G + 44:02:01G | 05:01:01G + 05:01:01G | 04:01:01G + 04:01:01G | 03:01:01G + 03:01:01G | 03:01:01i + 03:01:01i | 04:01:01i + 04:01:01i | NP + NP | 01:01:01i + 01:01:01i | NP + NP |
| OCILY3 | 6/12/2018 | 01:01:01G + 01:01:01G | 08:01:01G + 08:01:01G | 07:01:01G + 07:01:01G | 01:01:01G + 01:01:01G | 05:01:01G + 05:01:01G | 02:01:01i + 02:01:01i | 03:01:01i + 03:01:01i | 01:01:02i + 01:01:02i | NP + NP | NP + NP |
| OCILY8 | 8/03/2019 | 02:01:01G + 02:01:01G | * 51:01:01G + 58:01:01G | 03:02:01G + 14:02:01G | 02:02:01G + 05:01:01G | 03:01:01G + 05:01:01G | 02:01:01i + 03:03:02i | 03:01:01i + 09:01:02i | 02:02:01i + NP | 01:01:01i + NP | NP + NP |
| RIVA | 19/12/2018 | 02:01:01G + 03:01:01G | 44:02:01G + 47:01:01G | 05:01:01G + 06:02:01G | 04:01:01G + 04:01:01G | 01:03:01G + 05:01:01G | 02:01:01i + 06:03:01i | 03:01:01i + 13:01:01i | 01:01:02i + 01:01:02i | NP + NP | NP + NP |
| SC1 | 28/01/2019 | 02:01:01G + 02:01:01G | 44:02:01G + 44:02:01G | 05:01:01G + 05:01:01G | 04:01:01G + 04:01:01G | 01:02:01G + 01:02:01G | 06:02:01i + 06:02:01i | 15:01:01i + 15:01:01i | NP + NP | NP + NP | 01:01:01i + 01:01:01i |
| SUDHL4 | 28/02/2019 | 02:01:01G + 02:01:01G | 15:01:01G + 15:01:01G | 03:04:01G + 03:04:01G | 01:01:01G + 01:01:01G | 01:02:01G + 01:02:01G | 06:02:01i + 06:02:01i | 15:01:01i + 15:01:01i | NP + NP | NP + NP | 01:01:01i + 01:01:01i |
| SUDHL5 | 3/12/2018 | 03:01:01G + 24:02:01G | 07:02:01G + 52:01:01G | * 07:02:01G + 12:02:01G | 04:01:01G + 09:01:01 | 01:02:01G + 01:03:01G | 06:01:01i + 06:02:01i | 15:01:01i + 15:02:01i | NP + NP | NP + NP | 01:01:01i + 01:02i |
| TMD8 | 19/12/2018 | 02:07:01G + 02:07:01G | 46:01:01G + 46:01:01G | 01:02:01G + 01:02:01G | 05:01:01G + 05:01:01G | 03:01:01G + 03:01:01G | 03:03:02i + 03:03:02i | 09:01:02i + 09:01:02i | NP + NP | 01:01:01i + 01:01:01i | NP + NP |
| U2932 | 8/03/2019 | 01:01:01G + 03:01:01G | 08:01:01G + 15:01:01G | 03:04:01G + 07:01:01G | 03:01:01G + 04:01:01G | 03:01:01G + 05:01:01G | 02:01:01i + 03:02:01i | 03:01:01i + 04:01:01i | 01:01:02i + NP | 01:01:01i + NP | NP + NP |
| WSUDLCL2 | 8/03/2019 | 02:01:01G + 29:02:01G | 27:07:01G + 44:03:01G | 15:02:01G + 16:01:01G | 04:01:01G + 04:02:01G | 04:01:01G + 05:01:01G | 03:01:01i + 04:02:01i | 08:01:01i + 11:01:01i | 02:02:01i + NP | NP + NP | NP + NP |
| WSUFSCCL | 28/01/2019 | 02:01:01G + 02:01:01G | * 27:05:02G + 44:02:01G | 01:02:01G + 05:01:01G | 02:01:02G + 06:01:01G | 02:01:01G + 03:01:01G | 03:02:01i + 03:02:01i | 04:04:01 + 07:01:01i | NP + NP | 01:01:01i + 01:01:01i | NP + NP |

IMGT condensed ambiguous (v3320) HLA allele group results and the shaded cells (*) are putatively assigned base on CWD (v2.0) alleles; NP = allele not present;

*Karpas422 HLA-C
allele 1 - 07:02:01:17N or 07:02:01G or 07:02:01:17N
allele 2 - 07:02:01G or 07:02:01G or 07:02:01:17N
*OCILY8 HLA-B
allele 1 - 51:01:01G or 53:01:13
allele 2 - 58:01:01G or 58:08:02
*SUDHL5 HLA-C
allele 1 - 07:02:01G or 07:02:01:17N
allele 2 - 12:02:01G or 12:02:01G
*WSUFSCCL HLA-B
allele 1 - 27:127 or 27:05:02G
allele 2 - 47:04 or 44:02:01G

**Supplemental Figure 1. HLA typing of DLBCL tumor lines.** HLA typing of MHC-I and II loci of 18 DLBCLs commonly used throughout this work. NP, allele not present. Ambiguities listed at bottom.

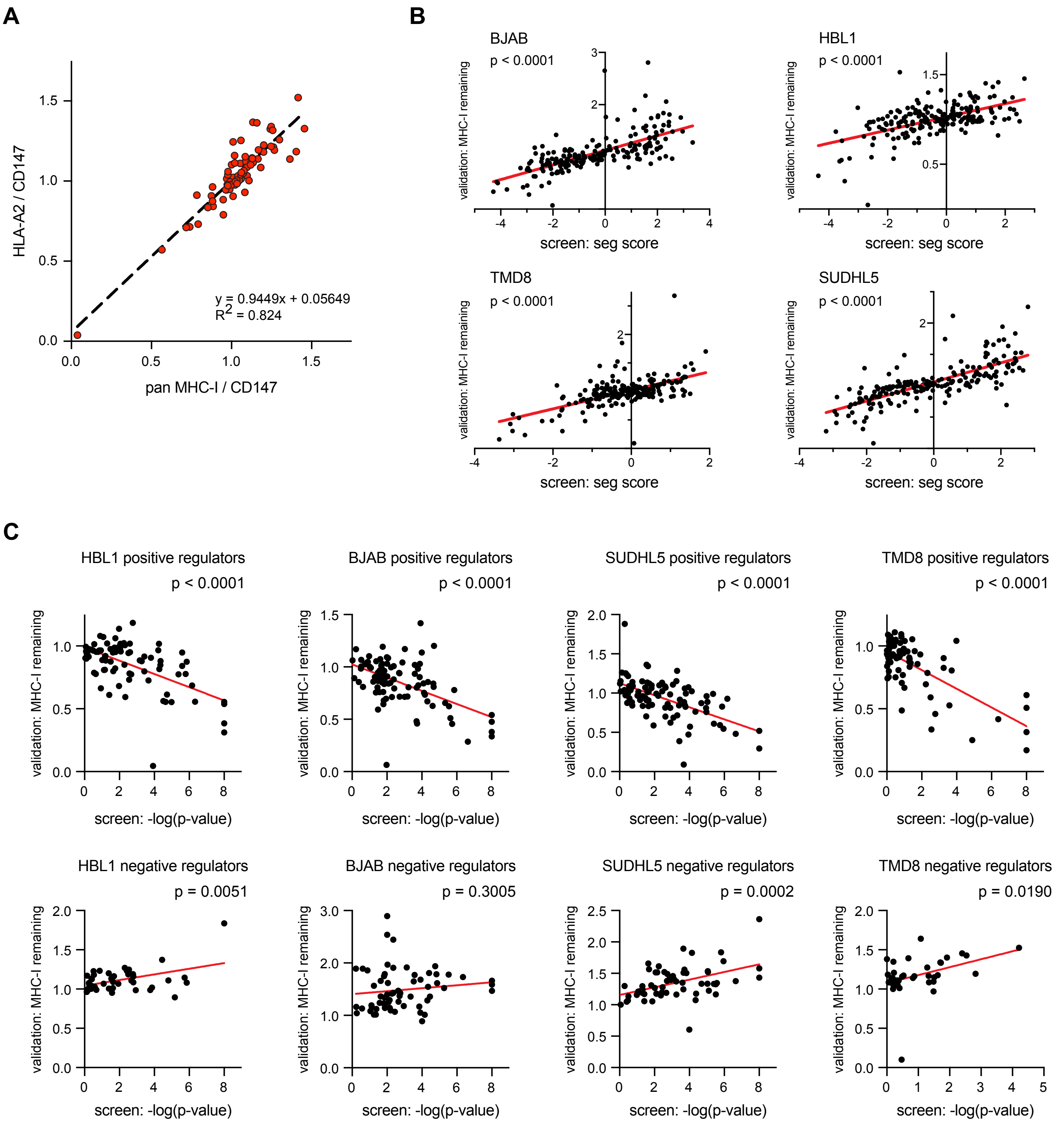


**Supplemental Figure 2. HLA-A2 correlation with pan-MHC-I measurements and screen correlation with KO cells. (A)** HLA-A2 per CD147 measurements were plotted against W6/32 per CD147 staining across a number of HBL1 KO cells; linear regression is plotted. **(B)** Individual KO cells were measured for surface MHC-I/CD147 and plotted against the CRISPR screen segregation score for the same gene. p-values represent likelihood of non-zero slope. **(C)** Surface MHC-I levels from individual gene KO cell lines during the validations were plotted against the -log(p-value) of the same gene from STARS analysis of the CRISPR screens. p-values represent likelihood of a non-zero slope. Only genes that were validated and received a score in STARS are plotted.

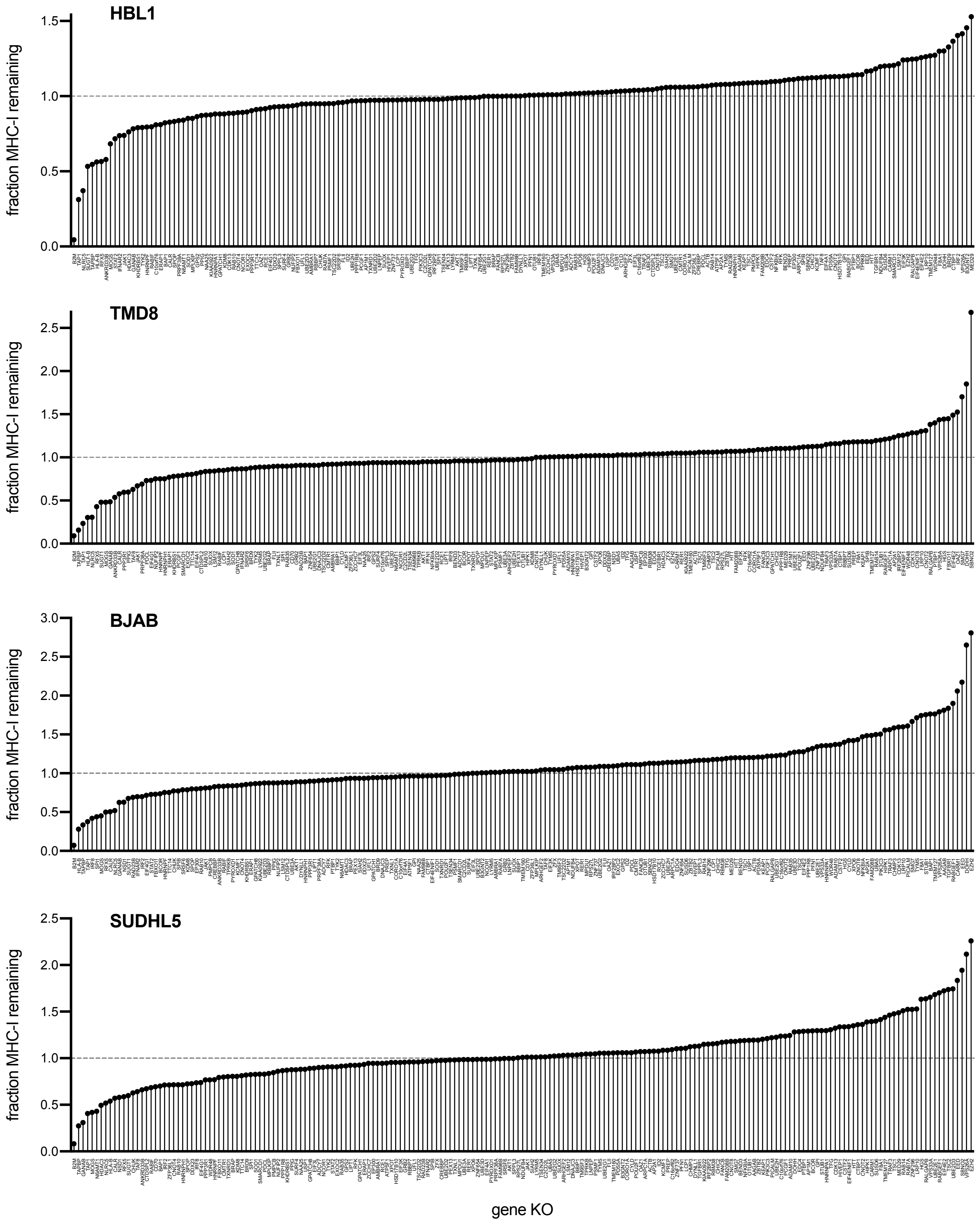


**Supplemental Figure 3. Summary of cell specific effects of gene KOs on MHC-I.** The fold change in surface MHC class I with KO of the indicated 199 genes, displayed across each cell model used for validation studies. See also Supplemental Table 3.

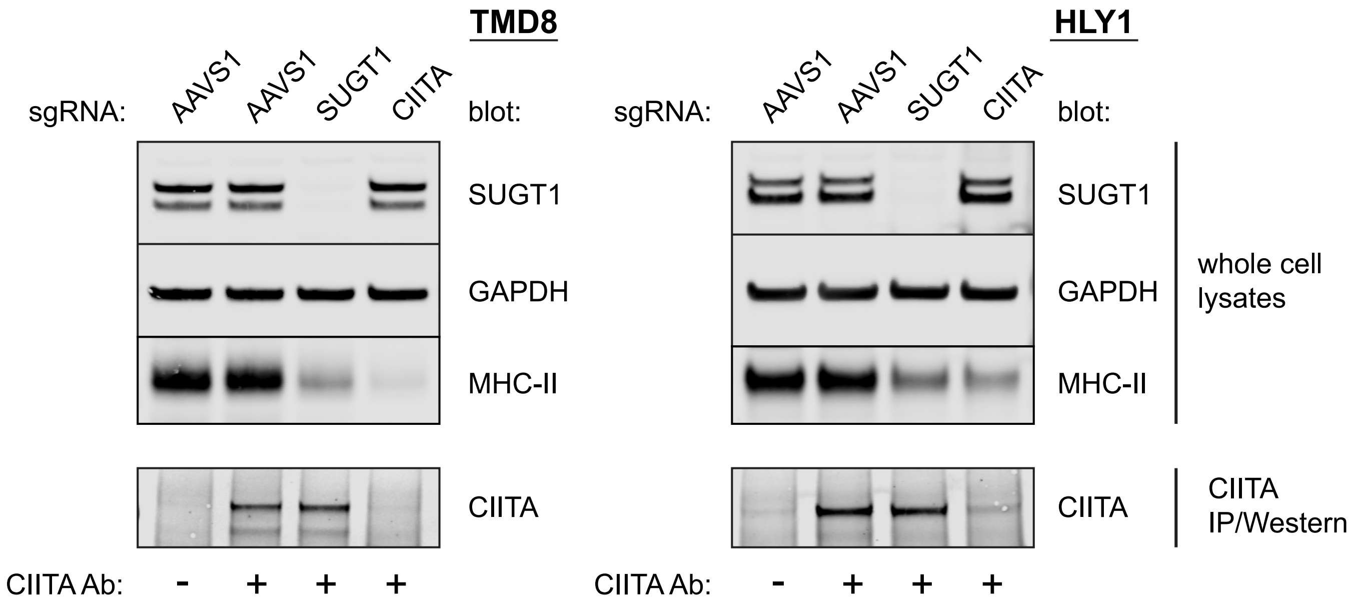


**Supplemental Figure 4. SUGT1 controls steady state levels of MHC-II.** TMD8 and HLY1 cells were infected with sgRNA against AAVS1 (negative control), SUGT1, or CIITA, and lysates were prepared for Western blotting. For detection of CIITA, IP/Western was required – independent antibodies were used for IP and blotting.

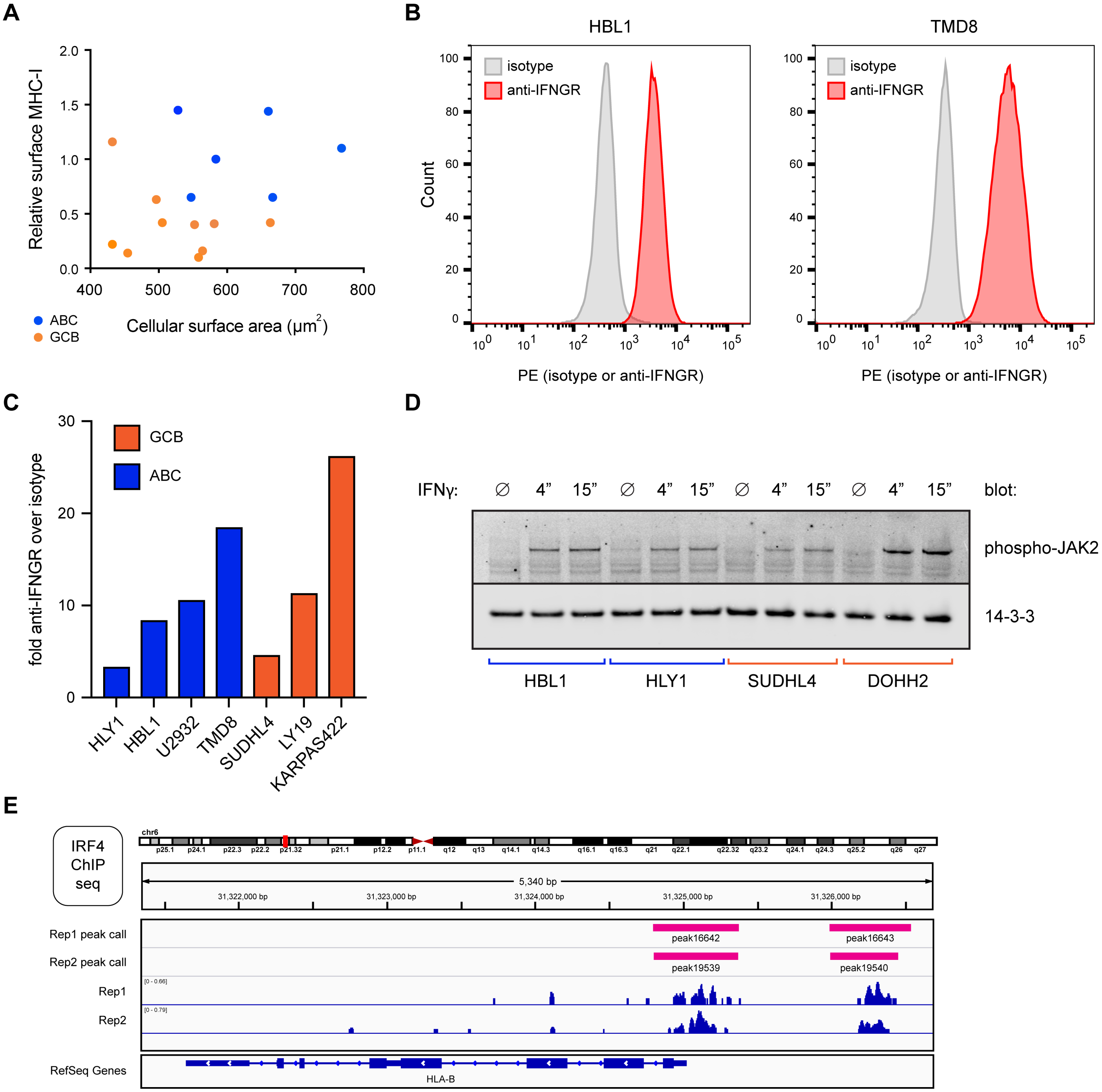


**Supplemental Figure 5. Extension of data related to Figure 5. (A)** Cell surface area was calculated by diameter measurements made by an automated cell counter and plotted against relative surface MHC-I levels (normalized to WT HBL1 cells). GCBs orange; ABCs blue. **(B)** Representative staining of IFNγ receptor is shown for HBL1 and TMD8 relative to isotype control. **(C)** A panel of 7 DLBCLs all show significant IFNγ receptor expression over isotype control staining, regardless of ABC/GCB classification. **(D)** The indicated cells were treated with 500U/mL of recombinant IFNγ for 4 minutes, 15 minutes, or left untreated prior to cell lysate preparation. Western blotting was performed with the indicated antibodies. **(E)** HLA-B gene and promoter region from published IRF4 ChIP-seq datasets of lymphoblastoid GM12878 (Gene Expression Omnibus, GSM803390).

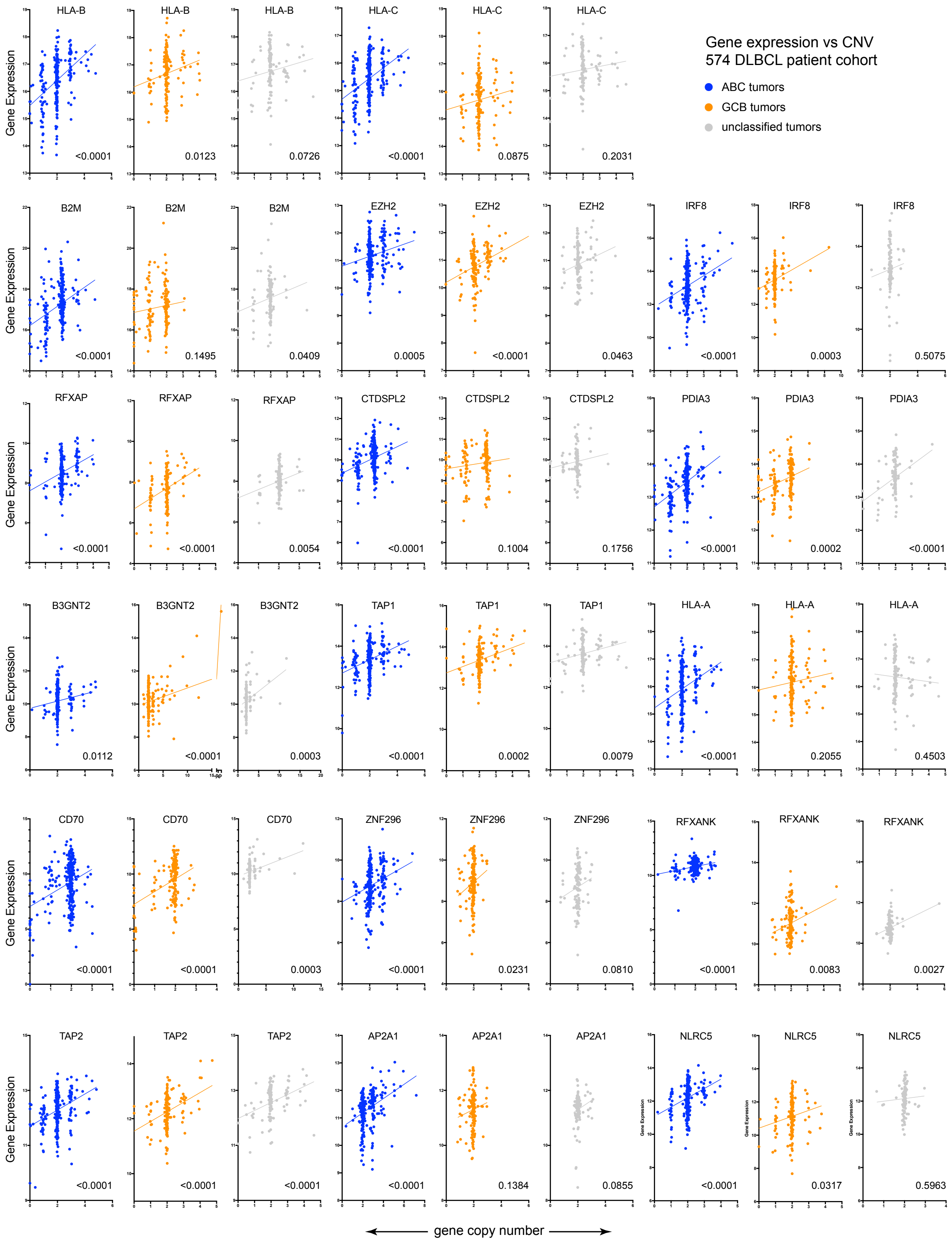


**Supplemental Figure 6. Copy number variation correlation with gene expression in DLBCL patient tumors.** Gene copy number variation is plotted versus gene expression for DLBCL patient tumor samples, indicating significant mRNA changes upon genetic copy number changes. p-values represent likelihood of positive correlation.

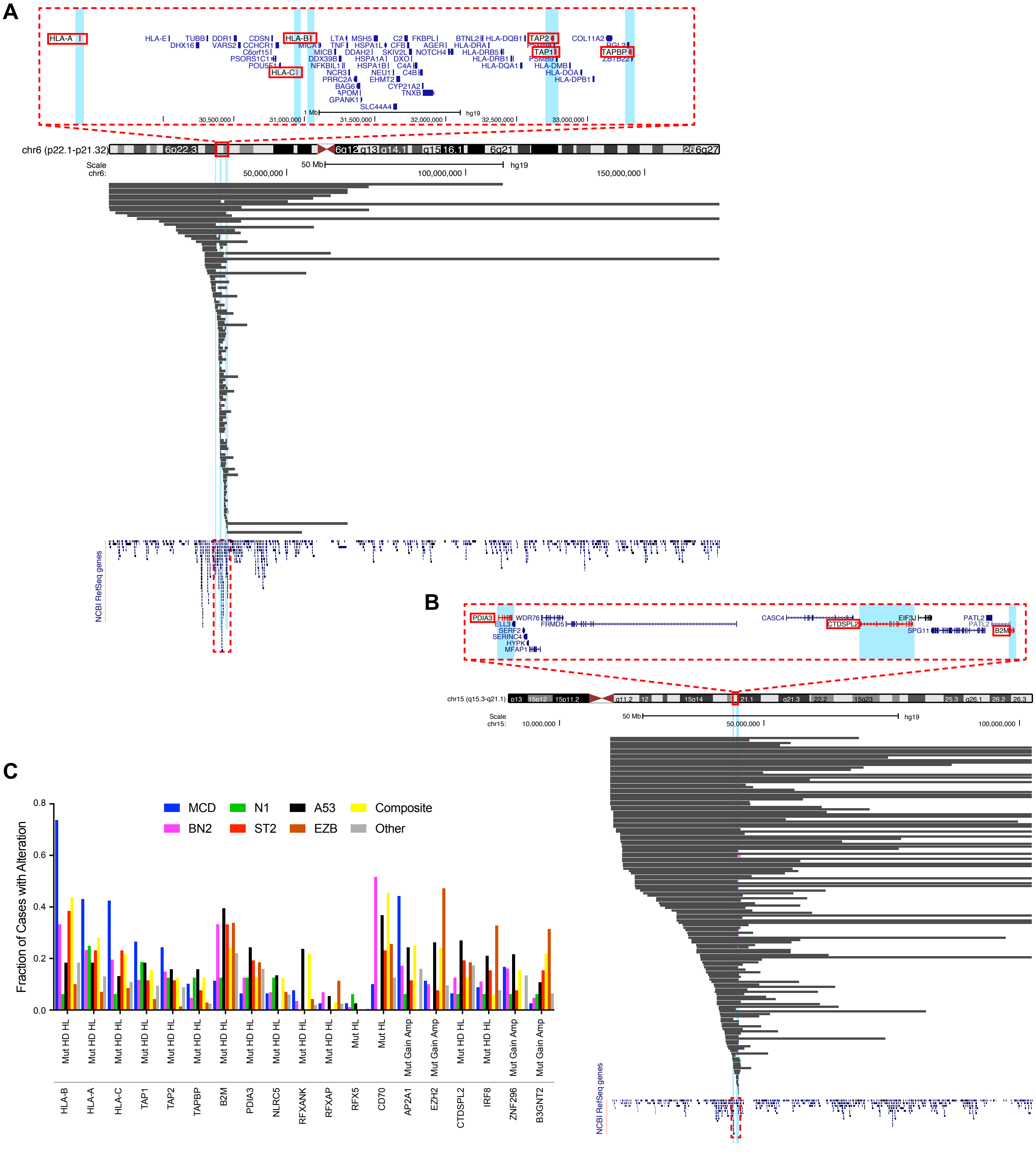


**Supplemental Figure 7. Patient chromosomal deletions and subtype classifications of mutations.** **(A)** Summary of chromosomal losses in patient tumors at the MHC locus, showing obvious selection of losses in APP machinery. **(B)** Same as **A**, but with chromosome 15. **(C)** Frequency of mutation in indicated genes across DLBCL patient cohort, sorted by LymphGen subclassification. Mut, mutation; HD, homozygous deletion; HL, heterozygous loss; gain, single copy gain; amp, multiple copy gain.

**A**

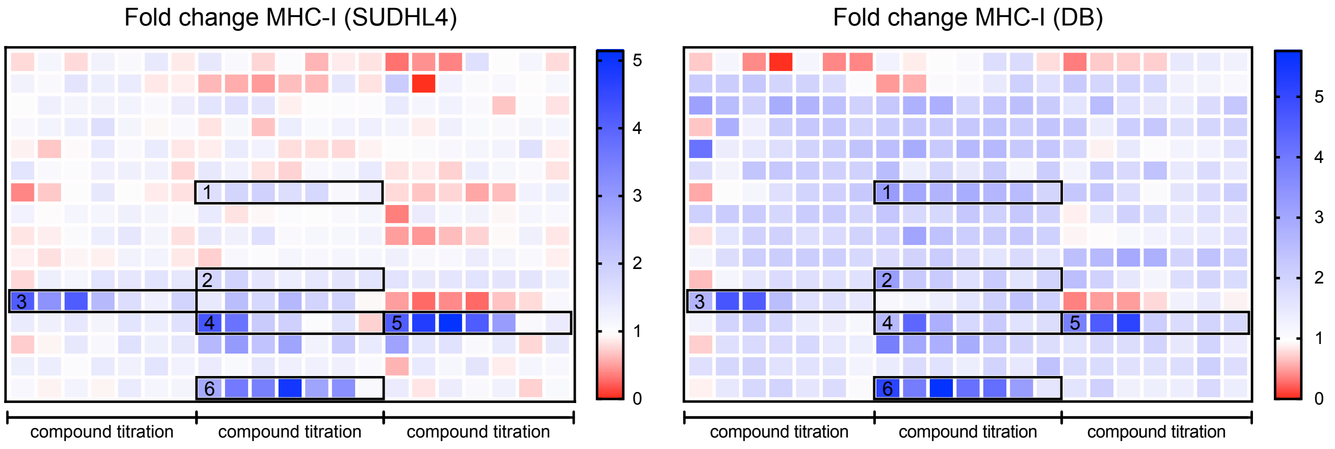


**B**

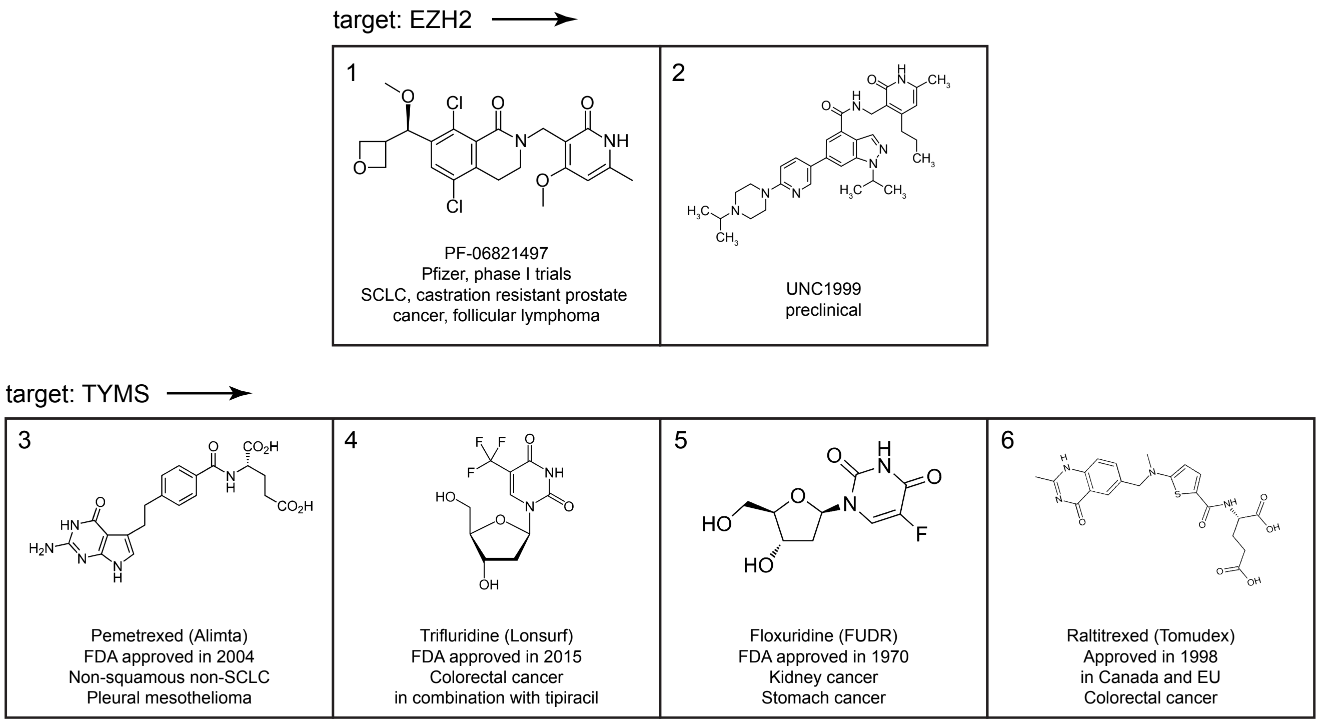


**Supplemental Figure 8. Targeted drug screen for inhibitors of negative MHC-I regulators. (A)** 48 selected small molecules were dry-spotted into wells of 384-well plates in a 7-dose titration series as indicated, spanning 5.7nM – 23.5 μM final concentrations upon cell plating. Cells were grown for 2 days prior to flow cytometry analysis of MHC-I surface expression. Fold change in surface MHC-I relative to vehicle controls is plotted by heatmap. Compounds 1-6, highlighted in black boxes, showed consistent, dose-dependent upregulation of MHC-I in both cell lines tested. **(B)** Chemical structures and clinical information related to compounds 1-6 above.

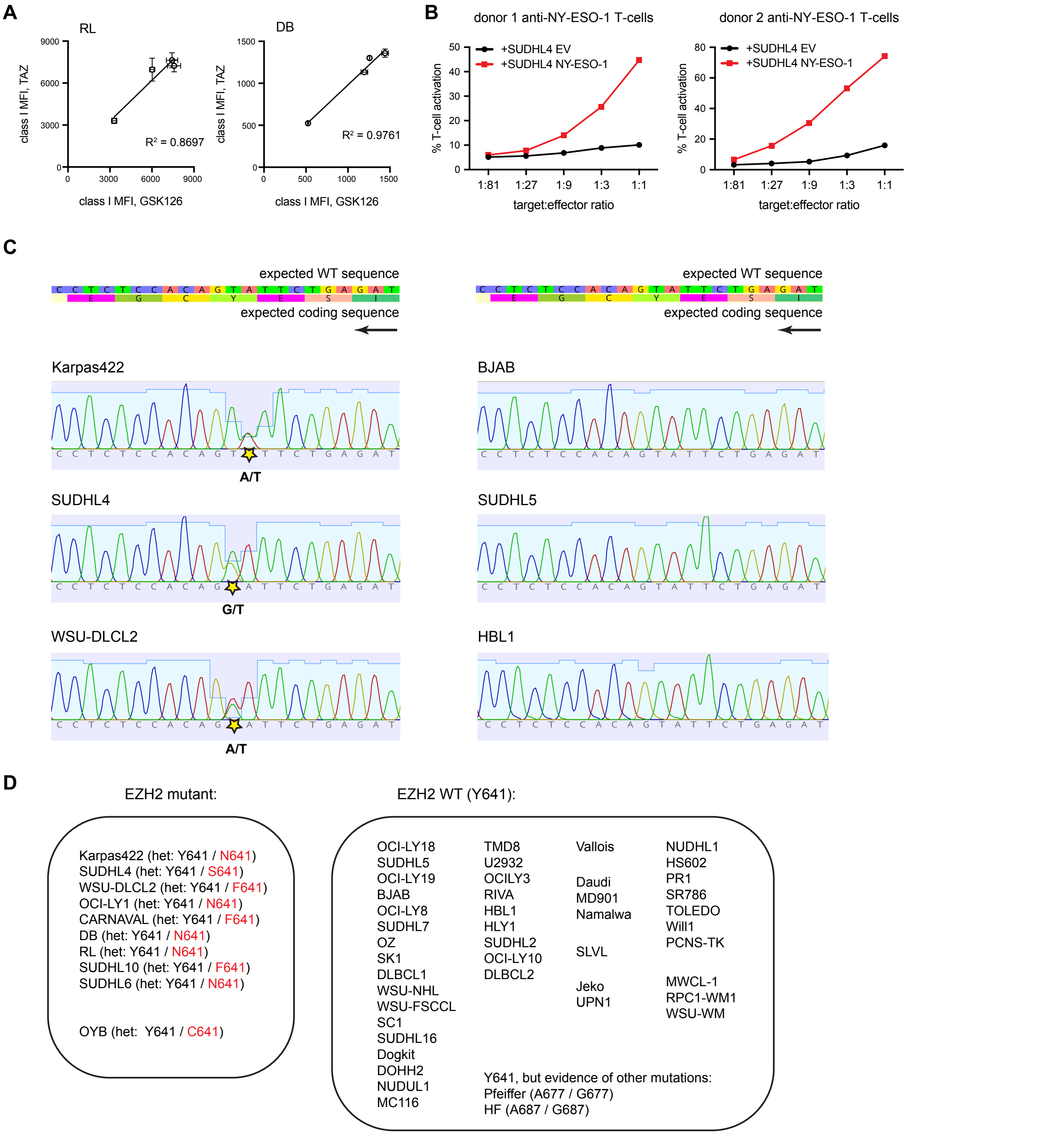


**Supplemental Figure 9. Extension of EZH2-related data in Figure 7. (A)** Treatment of DB or RL cells with increasing doses of tazemetostat or GSK126 for 7 days (DMSO, 0.4, 1.5, and 4μM). **(B)** Confirmation of NY-ESO-1 specificity in transduced SUDHL4 lines. T cells were stained for 4-1BB upregulation after co-culture with SUDHL4 cells transduced with either empty vector (EV) or NY-ESO-1 at the indicated target:effector ratios. **(C)** Example chromatograms of the EZH2 gDNA near Y641. Homozygous Y641 lines and heterozygous Y641 & Y641N/S/F are clearly distinguished. **(D)** Summary of EZH2 mutational status at Y641 across a number of tumor lines, with indicated heterozygous mutations in red.

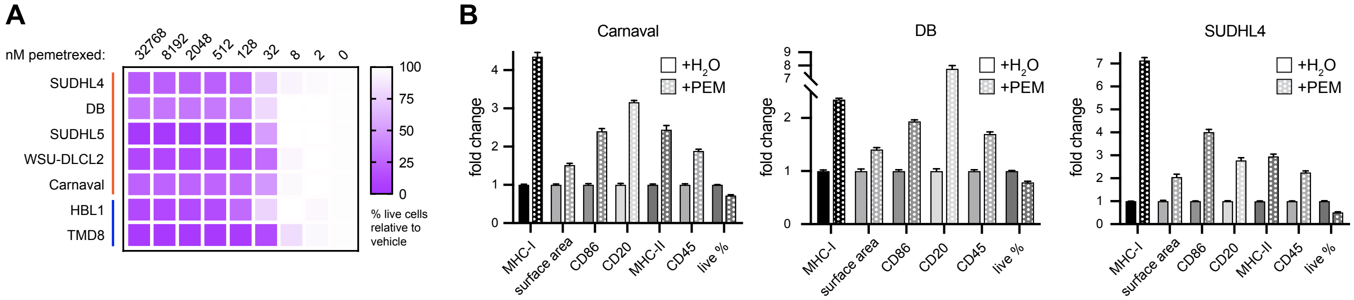


**Supplemental Figure 10. Extension of TS-related data in Figure 7. (A)** The indicated DLBCLs were cultured with serial dilutions of the TS inhibitor pemetrexed to determine % growth inhibition after 48 hours. **(B)** Cells were treated with pemetrexed or vehicle control and stained for various surface markers after 48 hours (Carnaval, 500nM; DB, 100nM; SUDHL4, 500nM). Surface areas were calculated by automated diameter measurements; "% live" indicated relative fraction of cells live by FACS scatter. For entire figure, bar graphs represent mean with standard deviations, minimum n = 3.
